# Structural basis for acyl chain control over glycosphingolipid sorting and vesicular trafficking

**DOI:** 10.1101/2021.04.26.440603

**Authors:** Stefanie S. Schmieder, Raju Tatituri, Michael Anderson, Kate Kelly, Wayne I. Lencer

## Abstract

The complex sphingolipids exhibit a diversity of ceramide acyl chain structures that influence their trafficking and intracellular distributions, but how the cell discerns among the different ceramides to affect such sorting remains unknown. To address mechanism, we synthesized a library of GM1 glycosphingolipids with naturally varied acyl chains and quantitatively assessed their sorting among different endocytic pathways. We found that a stretch of at least 14 saturated carbons extending from C1 at the water-bilayer interface dictated lysosomal sorting by exclusion from endosome sorting tubules. Sorting to the lysosome by the C14*-motif was cholesterol dependent. Perturbations of the C14*-motif by unsaturation enabled GM1 entry into endosomal sorting tubules of the recycling and retrograde pathways independently of cholesterol. Unsaturation occurring beyond the C14*-motif in very long acyl chains rescued lysosomal sorting. These results define a structural motif underlying membrane organization of sphingolipids and implicate cholesterol-sphingolipid nanodomain formation in sorting mechanisms.

## Introduction

Membrane function depends on the structural variations and dynamic physical properties of the acyl chains that form each class of its lipid components. But how the acyl chains operate to differentially affect the membrane biology of a lipid, and why so many different acyl chain structures are conserved within each lipid species remain in large part unexplained.

The complex sphingolipids, comprised of the glycosphingolipids (GSLs) and sphingomyelin, constitute 10-20 mole % of plasma membrane lipids (*1, 2*), and they provide a tractable experimental system to examine how acyl chain structure affects lipid function. In many cases, the activities of complex glycosphingolipids appear to depend upon locally segregating into liquid-liquid phase-separated nanodomains that form upon close interactions with cholesterol (*3–6*), the cortical cytoskeleton (*7–11*), and other scaffolding proteins (*12–14*). This is the lipid-raft or lipid nanodomain hypothesis (*15–20*). Considerable evidence shows that nanodomains are critical for key cellular processes including membrane protein trafficking, signal transduction, and receptor cell surface distributions (*16, 21–27*). Because of their functions in vesicular trafficking, some GSLs forming membrane nanodomains are co-opted by viruses and toxins, and by endogenous lectins, as platforms for entry into the cell and causation of disease (*28–34*).

Here, we studied ganglioside GM1, the glycosphingolipid receptor that mediates cholera toxin (CTx) entry into host cells. GM1 typifies the complex sphingolipids in basic structure and function. The different GM1 species, like all other complex sphingolipids, are comprised of a hydrophilic extracellular moiety (a specific pentasaccharide for GM1, one or more sugars for the other GSLs, and phosphocholine for sphingomyelin) coupled to a ceramide that anchors the molecule in the membrane. The ceramide moieties consist of a sphingosine base (predominantly 18 hydrocarbons long, C18:1^Δ4^, *trans*) linked to a fatty acid by an amide bond to the C1 carbon positioned at the lipid bilayer-water interface. The sphingosine base varies in structure only to a small degree, but the acyl chain can display considerable heterogeneity in both length (C14 to C26) and saturation (single *cis* or no double bond) (*35, 36*). Because these lipids contain large hydrophilic extracellular head groups, they are trapped in the outer leaflet of the membrane bilayer and their distribution throughout the cell occurs only by vesicular traffic (*37, 38*). This can be quantitatively measured and compared for GM1 species with identical head groups but with varied acyl chains in their ceramide moieties.

We previously found the different acyl chains in the ceramide moiety of GM1 act decisively in trafficking the toxin into the different endocytic and sub-cellular compartments required for the induction of toxicity (*39, 40*). GM1 species containing ceramides with “kinked” shapes caused by a *cis*-double bond in the acyl chains (C16:1^Δ9^ or C18:1^Δ9^) sorted into the recycling and retrograde endosome pathways, and GM1 species containing ceramides with saturated acyl chains of the same length, C16:0 and C18:0, but with “straight conical” shapes, sorted to the lysosome instead (*40*). Additional evidence that acyl chain structure of the complex sphingolipids may affect trafficking was recently shown for ceramide-anchored proteins in the biosynthetic pathway of yeast (*41*). But how the cell discerns among the different ceramide moieties to affect this acyl chain-based sorting has not been explained.

To address these problems, we synthesized a library of eleven GM1 species with ceramides reflecting the naturally occurring variations of fatty acyl chain length and saturation. This provided a direct test of GSL structural biology in ways not previously possible.

In mammals, the single *cis*-double bond in the acyl chain of the unsaturated ceramide is created by palmityl/stearoyl-CoA desaturase (SCD). The enzyme installs a *cis*-double bond invariably at the Δ^9^-position of C16:0 or C18:0 fatty acids, creating a ‘kinked’ shaped lipid (*42*). When coupled to sphingosine by the ceramide synthases, and then incorporated into the complex sphingolipids, the kink in the acyl chain of these ceramide species is positioned where it should impair close packing against the rigid planar surface of cholesterol (*43*). On the other hand, GSL species containing ceramides with longer unsaturated acyl chains will have the *cis*-double bonds placed deeper within the membrane bilayer. They are built upon the same C18:1 acyl chains generated by the Δ9-desaturase, but lengthened at the carboxy-terminus by the ELOVL elongases (*44*). Consequently these unsaturated very long acyl chains contain longer stretches of saturated hydrocarbons extending from the water-bilayer interface into the membrane bilayer. In some cases, the *cis*-double bond will be positioned beyond the planar surface of cholesterol. We reasoned that these unsaturated, and “kinked”, very long acyl chain GSLs might still assemble closely with cholesterol to form nanodomains and function accordingly - thus providing a structural test of sphingolipid behavior in live cells.

Our results define a structural motif in the ceramide acyl chain of the GSLs that dictates endosomal sorting and with broad implications for current models of membrane structure and function.

## Results

The eleven GM1 molecules synthesized for testing contained saturated and mono-unsaturated acyl chains (Fig. 1B) produced naturally in mammalian cells (*45, 46*). Three additional molecules, the GM1 C12:1^Δ6^, C12:0, and C16:1^Δ9^ species, were prepared to contain non-native ceramide acyl chains that broadened the testing of the hypothesis (Fig. 1B, Suppl. Fig. 1A). The cellular and membrane behaviors of each GM1 species were studied by coupling the extracellular oligosaccharide to a ‘reporter’-peptide containing biotin and fluorophore (Suppl. Fig. 1A), or directly to a fluorophore (*40, 47, 48*).

**Fig. 1.**
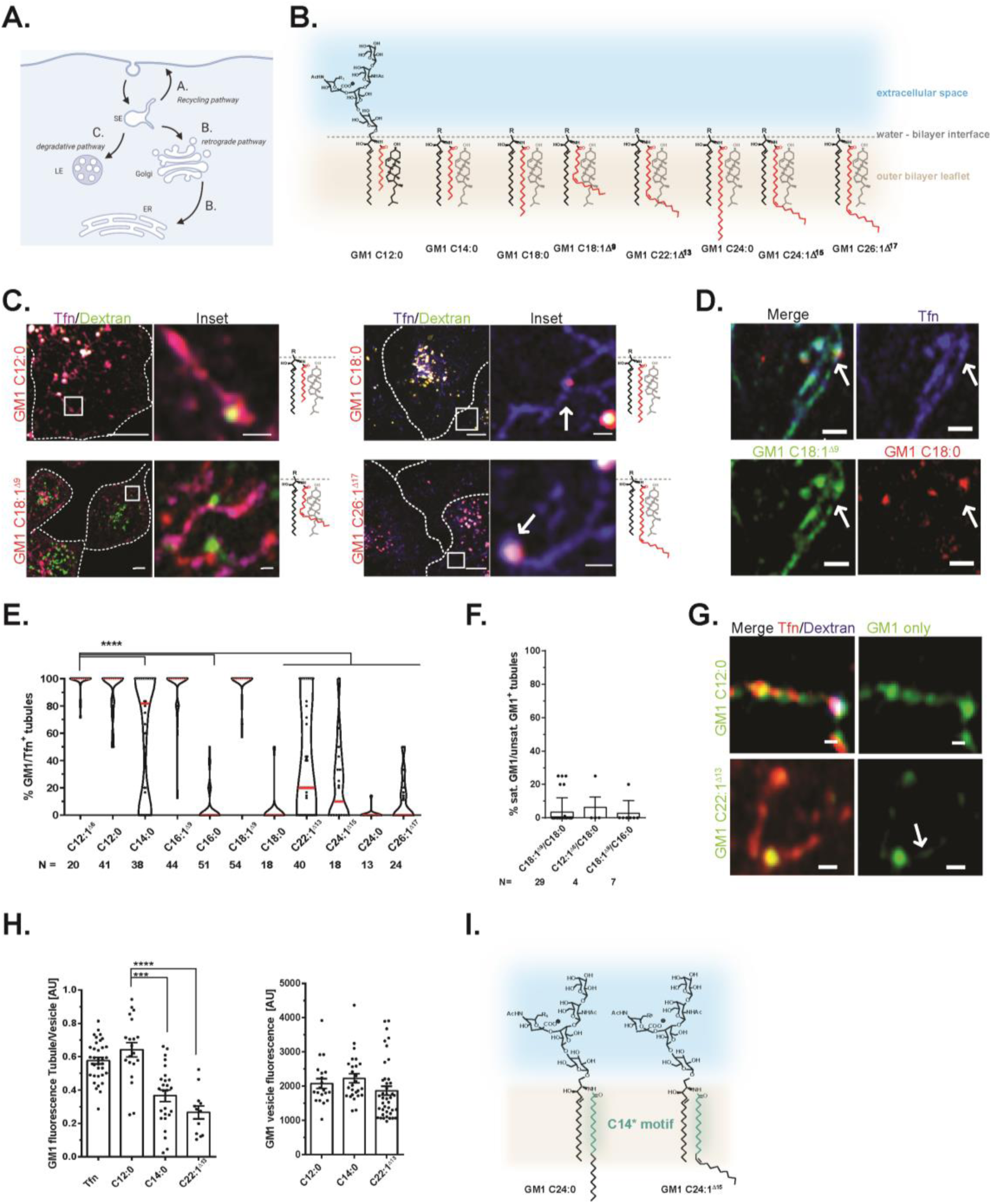
Ceramide structure-specific trafficking within the sorting endosome. **(A)** Schematic of endocytic pathways emerging from the sorting endosome (SE). Both the recycling and retrograde pathways (marked A and B) require sorting into narrow endosomal tubules. Cargos destined for degradation are retained within the sorting endosomal body for maturation to late endosomes (LE) (marked C). **(B)** Library of GM1 species synthesized - acyl chains marked in red and arbitrarily aligned against cholesterol to mark changes in structure. GM1 C12:1^Δ6^, C16:0 and GM1 C16:1^Δ9^, species not shown. **(C)** Airyscan live cell imaging of recycling endosomal vesicles in A431 epithelial cells. Scale bars = 5 and 0.5 μm respectively. **(D)** Same as panel C for cells treated with GM1 C18:1^Δ9^ (green) and GM1 C18:0 (red). Scale bar = 1 μm. **(E)** Quantification of GM1 presence in Tfn marked recycling endosomal tubules. Mean ± SEM, **** p = < 0.0001 by one-way ANOVA and Tukey’s multiple comparison using GM1 C12:0 as comparison. **(F)** Co-localization of GM1 C18:0 or C16:0 with GM1 C18:1^Δ9^ or C12:1^Δ6^ species in recycling Tfn-positive endosomal tubules. Quantification and statistics as in panel E. **(G)** Recycling tubules analyzed as in Panel C using GM1 C22:1^Δ13^ and GM1 C12:0 species (green), Tfn (red). Scale bar = 0.5 μm. **(H)** Left panel: Quantification of mean grey values of GM1 fluorescence in Tfn tubule normalized to GM1 fluorescence in body of recycling endosomes for GM1 C12:0, C14:0 and C22:1 ^Δ13^. Right panel: Mean grey value of GM1 fluorescence in body of recycling endosome for the same GM1 species. Mean ± SEM. **** p = < 0.0001 against GM1 C12:0 by One-way ANOVA and Tukey’s multiple comparison. **(I)** GM1 C24:0 and GM1 C24:1^Δ15^, C14* motif in turquoise.

As a first test of membrane behavior, the liquid ordered (L_o_) and liquid disordered (L_d_) phase preferences for each of the GM1 species were defined *in vitro* using giant plasma membrane vesicles (GPMVs) prepared from different cell lines as indicated (Suppl. Figs. 1B – D). Certain saturated lipids and sterols can condense into higher ordered assemblies within membranes, termed the L_o_ phase. This separates them from the other lipids present in the same membrane termed the L_d_ phase. Phase partitioning into the L_o_ phase can be considered a feature of nanodomains in live cells (*49, 50*). We found the major determinant for the phase behaviors of the different GM1 species in GMPVs to be the presence or absence of a *cis*-double bond in the ceramide acyl chain. The unsaturated GM1 species segregated into the L_d_ phase irrespective of the cell type-specific GMPV tested. GM1 with fully saturated ceramide acyl chains associated exclusively with the L_o_ phase. Thus, partitioning into the L_o_ phase in GPMVs depended on full saturation of the fatty acyl chain.

### A saturated acyl chain length of at least 14 carbon atoms in the GM1 ceramide directs sorting to the lysosome

We next used lattice light-sheet and airyscan microscopy to image the different fluorescently-labeled GM1 species in the sorting endosome and their ability to enter endosomal sorting tubules. Endosomal sorting tubules affect the trafficking of membrane and cargo away from the lysosome and into the recycling pathway for return back to the plasma membrane (PM), or into the retrograde pathway for transport to Golgi and endoplasmic reticulum (ER) (schematic, Fig. 1A). For these studies, human A431 epithelial cells were co-incubated with nM concentrations of the different GM1 species together with transferrin (Tfn) and dextran. Tfn and dextran were used to mark the endosome recycling and the lysosome pathways respectively (*51, 52*). Sorting endosomes, which serve all pathways, were operationally defined as Tfn and dextran double positive vesicles, and sorting tubules emerging from these structures were identified by their Tfn-only content. We tested several cell lines for this assay. The A431 cell line selected for these studies exhibited highly elongated and easily visualized tubules emerging from the sorting endosome. The same basic phenotypes, however, were observed in other A431 clones, and HeLa (Suppl. Fig 1D and E), MDCK II, Caco2, and HT29 cell lines which all lacked the unusually elongated tubules (data not shown).

We found that the GM1 species with ceramides containing short saturated acyl chains (≤ C14:0) or unsaturated acyl chains C12:1^Δ6^ through C22:1^Δ13^ efficiently co-localized with Tfn in endosome sorting tubules (Fig. 1C, left panels and quantified in Fig. 1E and Suppl. Video 1). The GM1 species with ceramides containing long saturated acyl chains ≥ C16:0, however, did not (Fig. 1C, right panels, Fig. 1E and Suppl. Video 2). They were localized only to the body of the sorting endosome. We tested this observation again using the C18:0 and C18:1 ceramide pair loaded together into the same cells and obtained the same results (Fig. 1D and 1F and Suppl. Video 2).

Notably, we also found that the GM1 species containing ceramides with unsaturated but very long acyl chains (≥ C24:1^Δ15^ in length) were localized only in the body of the sorting endosome, behaving like the GM1 species with fully saturated acyl chains. These GSLs did not enter the narrow highly-curved sorting tubules despite the presence of a *cis*-double bond in their ceramide acyl chain (Fig. 1B, 1C lower right panels, and Fig. 1E). Additionally, the GM1 species with C14:0 or C22:1^Δ13^ fatty acids displayed intermediate and closely similar phenotypes. They still entered endosome sorting tubules but less efficiently (measured as lower fluorescence intensity in recycling endosomal tubules) (Fig. 1G and H left panel). This observation was not explained by reduced endocytic uptake of the lipids (Fig. 1H right panel). Closely similar phenotypes for all the GM1 species tested were found in Hela cells (Suppl. Fig. 1E and F), and for the other epithelial cell lines noted above (data not shown). Thus, while the presence or absence of a *cis*-double bond in the ceramide acyl chain dictated L_o_ and L_d_ phase partitioning in GMPVs, the position of the double bond within the acyl chain acted as the decisive factor for endosomal sorting in live cells. Lipid shape, on its own, was not sufficient to explain the differential sorting observed. These results, especially considering the intermediate behaviors of the C14:0 and C22:1^Δ13^ ceramide species, showed that a saturated acyl chain length of at least 14 carbon atoms plus one, extending from the C1 amide bond at the water-bilayer interface, was necessary for retention of GM1 in the body of the endosome. We henceforth name this structure the C14*-motif (Fig. 1G).

### Independence from Lectin binding

The C14*-motif may be necessary but not sufficient to explain the differential endosomal sorting of GM1 as endogenous lectins capable of scaffolding the lipid may be involved (*34*). Such an effect on GM1 trafficking is perhaps best typified by multi-valent binding to CTxB (*33, 40, 53-55*). As such, we first tested if binding CTxB might act on the different GM1 species to affect changes in their L_d_ and L_o_ phase behaviors as assessed in GMPVs. The addition of CTxB caused a switch from L_d_ to L_o_ phase preference, but only for the GM1 species with very long unsaturated acyl chains ≥ 22:1^Δ13^ (Suppl. Fig.1G). The phase preferences for the other GM1 species were not affected.

To test if in live cells cross-linking GM1 by endogenous lectins may be required for its endosomal sorting, we studied the GM1 C16:0 and C16:1^Δ9^ pair, and GM1 C24:1^Δ15^. Here, intracellular trafficking of the different GM1 species was studied in A431 cells continuously exposed to 100 mM lactose or sucrose to compete against binding the GM1 oligosaccharide moiety by endogenous lectins (*56*). We found no effect of the disaccharide treatments (Suppl. Fig. 1H). Control studies showed that 100 mM lactose was sufficient to block the high affinity binding of CTxB to GM1 on the A431 cell surface (Suppl. Fig.1I). Thus, binding by endogenous galactose- or glucose-binding lectins does not appear to be required for the observed GM1 endosomal sorting (Fig. 1B) - in these cases, the C14*-motif is both necessary and sufficient.

### Differential sorting between the lysosomal and recycling pathways

To quantitatively test if endosomal GM1 sorting is based on the C14* motif, we developed a FACS based assay to measure GM1 sorting between recycling endosomes and lysosomes (Fig. 1A, Pathways A vs C). For these studies, we synthesized the GM1 library coupled to the all-D amino acid ‘reporter’-peptide containing only a biotin functional group (Suppl. Fig. 1A). Conditions for incorporating the different GM1 species into the PM at similar amounts were determined experimentally (Fig. 2A, left panel). After a 3h ‘chase’, the amount of the biotin-labeled-GM1 located at the cell surface was determined by FACS measured by binding of fluorescently-labeled streptavidin (Fig. 2A and 2B). We found that the GM1 species which entered endosome sorting tubules and lacked the C14* motif (GM1 C12:0, C14:0, C16:1^Δ9^, C18:1^Δ9^ and C22:1^Δ13^, Figs. 1E and 2A and B magenta points) were recycled after endocytosis. They were detected on the PM at significantly higher levels compared to the GM1 species containing the C14*-motif (GM1 C16:0, C18:0, C24:1^Δ15^, C24:0 and C26:1^Δ17^, Fig. 2A and 2B turquoise points). Even after longer chase times (> 6 h), the GM1 C16:1^Δ9^ species still localized to the PM and within endosomal recycling tubules containing Tfn; while the GM1 C16:0 species localized strictly to intracellular vesicles stained by lysotracker™ (marking late endosomes and lysosomes, Suppl. Fig. 2A). We observed no confounding effects of GM1 treatment on endocytosis per se - as measured by dextran (fluid-phase) or Tfn (receptor-mediated) uptake (Suppl. Fig. 2B). In addition, the results could not be simply explained by differential endocytic uptake mechanisms into caveolae - as neither the C18:1^Δ9^ nor C18:0 species were sensitive to CRISPR-Cas9 deletion of cavin1 (Suppl. 2C). Thus, a saturated acyl chain length of at least 14 saturated carbon atoms plus one (the C14*-motif) drives endosomal sorting of GM1 to the lysosome, with GM1 C14:0 and C22:1^Δ13^ displaying intermediate phenotypes (Fig. 1A pathway C, and Fig. 2C). We note again that lysosomal sorting was rescued for the unsaturated GM1 species containing very long unsaturated acyl chains (i.e. unsaturated “kinked shaped” fatty acids containing the C14*-motif”).

**Fig. 2.**
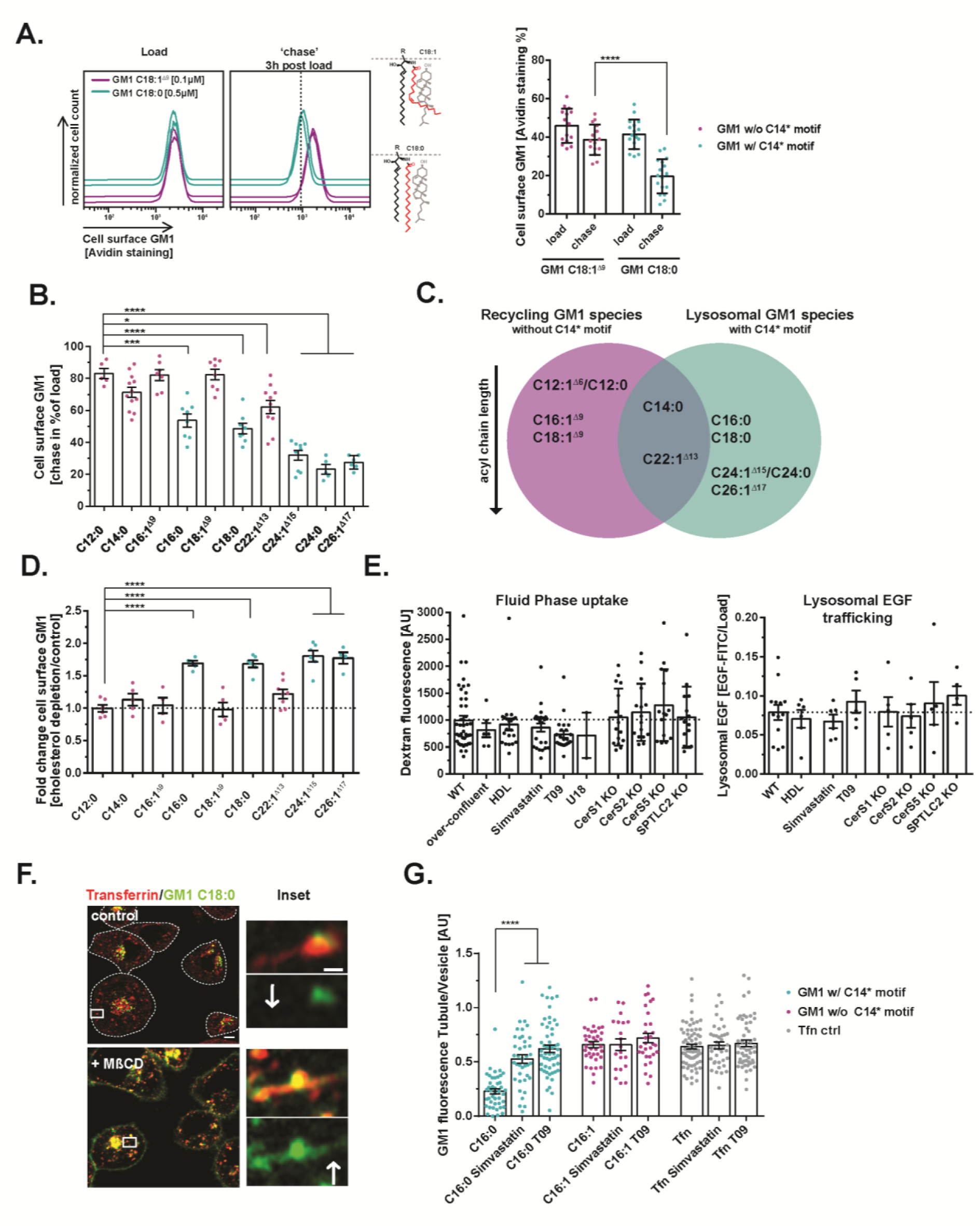
C14* motif dictates differential sorting between the lysosomal and recycling pathways. **(A)** PM concentrations of GM1 C18:0 (turquoise, with C14* motif) and GM1 C18:1^Δ9^ (magenta, without C14* motif) as assessed by FACS using AlexaFluor™-labeled streptavidin before and after 3 h chase. Mean ± SEM (right panel), each dot equals 2 biological replicates measuring 20,000 cells each. **** p = < 0.0001 by unpaired t-test. **(B)** Plasma membrane recycling as in panel A for the GM1 structural library (GM1 species with C14* motif in turquoise, GM1 species without C14* motif in magenta). Mean ± SEM. One-way ANOVA and Tukey’s multiple comparison. **(C)** Venn diagram for ceramide acyl chain structure dependent GM1 sorting in the endosomal system. Recycling GM1 species, without C14* motif are in magenta and GM1 species entering the lysosomal pathway, containing the C14* motif are in turquoise. **(D)** Change in recycling rates under mildly cholesterol depleting conditions as indicated. **** p = < 0.0001 by One-way ANOVA and Tukey’s multiple comparison. **(E)** Fluid phase uptake (left) and lysosomal trafficking (right) measured by FACS in A431 cells depleted or not of cholesterol as indicated. No differences were observed as measured by One-way ANOVA and Tukey’s multiple comparison. **(F)** Images of recycling tubules in A431 cells treated or not with MβCD. Scale bars = and 0.5 μm respectively. **(G)** Entry of GM1 C16:0 or C16:1^Δ9^ into recycling tubules in cholesterol depleted A431 cells as indicated. Mean grey values of fluorescent GM1 in both endosomal vesicle as well as tubule was recorded and the ratio of GM1 fluorescence in tubule over vesicle was calculated for each condition. Each dot represents one tubule vs vesicle ratio. Mean ± SEM. **** p = < 0.0001 by One-way ANOVA and Tukey’s multiple comparison.

### Cholesterol Dependence for Sorting to the Lysosome

The nanodomain (lipid raft) hypothesis posits strong dependence on close assembly between GSLs and membrane cholesterol. We reasoned, however, that such cholesterol dependence should be evident for endosomal sorting of only the GM1 species with ceramide structures containing the C14*-motif - if the motif acts as we hypothesize by allowing close alignment against the rigid and flat surface of cholesterol (*3, 6, 43*). Likewise, endosomal sorting of the GM1 species with *cis*-double bonds at positions disrupting the C14*-motif should behave independently of membrane cholesterol. But the GM1 species with a *cis*-double bond located beyond the C14*-motif, as seen in the naturally occurring very long acyl chains (> C22:1^Δ13^ acyl chains), should again show cholesterol-dependence.

To test this idea, we studied the endosome recycling of the different GM1 species in A431 cells mildly reduced of membrane cholesterol: either by treatment with low doses of the small molecule inhibitors of cholesterol metabolism simvastatin, T0901317, or U18666A; or as we found experimentally by CRISPR-Cas9 knock out of serine-palmityltransferase-2 (SPTLC-2), ceramide synthase 5 (CerS5), or ceramide synthase 2 (CerS2), or by simply growing the cells in over-confluent conditions. In all cases, the experimental conditions were found to lower plasma membrane cholesterol content as measured by loss of filipin staining (Suppl. Fig. 2D, left panel), and further evidenced by increase in the general polarization (GP) of laurdan as assessed in GPMVs (implicating increased membrane fluidity typifying cholesterol depleted membranes, Suppl. Fig. 2D, right panel). We also conducted lipidomic profiling by mass spectrometry, which showed in all cases that cells depleted of cholesterol had only minor changes in the different species of the major membrane lipid classes: phosphatidylethanolamine (PE), phosphatidylglycerol (PG), and phosphatidylcholine (PC) (Suppl. Fig. 3D). Additionally, and in all cases, cells depleted in cholesterol by our methods retained their normal endosomal and overall morphologies and endocytic uptake rates with only minor variations (Fig. 2E left panel and Suppl. Fig. 2E). We also note that cholesterol depletion induced by over-confluent cell culture, the least invasive perturbation, had only minimal effects on the overall lipidome and no detectable changes on phase preferences for any of the different GM1 species in GPMVs (Suppl. Fig. 2H).

Consistently, for all methods, we found no effect of cholesterol depletion on the recycling of GM1 species lacking the C14*-motif (Fig. 2D magenta points, and Suppl. Fig. 2F, left table). In contrast, cholesterol depletion uniformly affected the late endosome/lysosome sorting of GM1 species with ceramide moieties containing the C14*-motif. This included the saturated acyl chains ≥ C16:0 and notably the unsaturated ceramides with very long unsaturated acyl chains ≥ C24:1^Δ15^. In cholesterol depleted cells, these GM1 species were no longer strictly sorted into the late endosome/lysosome - instead, large fractions of these lipids were recycled back to the PM (Fig. 2D turquoise points and Suppl. Fig. 2F, right table) and co-localized within Tfn-containing endosome recycling tubules (where none was found before) (Fig. 2F and G). The effect of cholesterol depletion on lysosomal sorting was specific for GM1, as lysosomal sorting for the epidermal growth factor (EGF) receptor or the low density lipoprotein receptor LDL-R were not affected (Fig. 2E right panel and Suppl. Fig. 2G). Thus, the sorting step for trafficking GM1 into the lysosome appears to be dependent on membrane cholesterol, but only for the GM1 species with ceramide structures containing the C14*-motif and predicted to interface tightly alongside the sterol - consistent with the nanodomain hypothesis.

### Sorting into retrograde and recycling pathways by bulk membrane flow

We next considered the possibility that the GM1 species lacking the C14*-motif may enter endosome sorting tubules by default, following bulk membrane flow (*57*). If this were the case, we reasoned these GM1 species should enter all pathways emerging from the sorting endosome, including the retrograde pathway to the trans Golgi and ER. For these studies, we used the B-subunit of cholera toxin (CTxB). Though CTxB scaffolds GM1 into a multimeric complex affecting its membrane behavior (*54, 58, 59*), ceramide structure still plays a decisive role in endosomal sorting into the ER (*40*) and in liquid phase partitioning in GMPVs (Suppl. Fig. 1G). For these studies, the different GM1 species were applied to a HeLa cell line that endogenously lacks GM1. CTxB trafficking into the ER was quantitatively measured using a FACS-based split-GFP assay recently developed by our group (*60*). We found that only the short chain (≤ C14:0) or unsaturated GM1 species (≤ C22:1^Δ13^), both lacking the C14*-motif, efficiently trafficked CTxB into the ER (Fig. 3A and 3B). The GM1 species containing the C14*-motif did not. This includes the GM1 species with ceramide structures containing long saturated acyl chains (≥ C16:0), and those containing unsaturated but very long acyl chains (≥ C24:1^Δ15^), which rescues the C14*-motif and its function. Thus, the requirement for acyl chain structure directing the different GM1 species into the retrograde endosomal pathway phenocopied exactly the structures required for sorting into the recycling pathway.

**Fig. 3.**
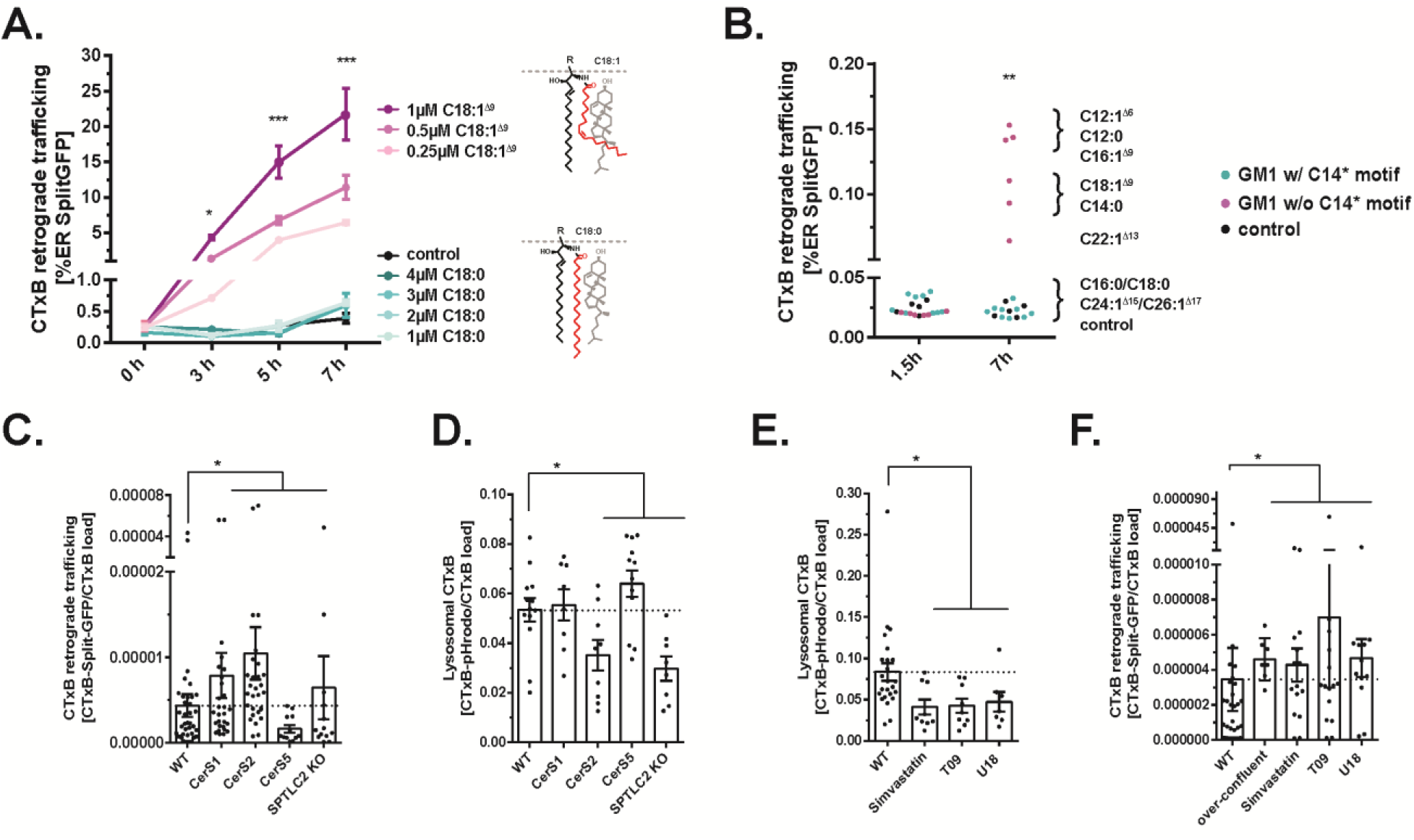
Ceramide-dependent sorting into the retrograde pathway. **(A)** Time course of retrograde GM1 trafficking to the ER by a SplitGFP based FACS assay Graph dots represent the mean ± SD of 2 biological replicates measuring 20,000 cells each. **** p = < 0.0001 by One-way ANOVA and Tukey’s multiple comparison. **(B)** SplitGFP assay as in (A) loaded with different GM1 species. Graph dots represent the mean ± SEM of 2 biological replicates measuring 20,000 cells each. **(C)** A431 CRISPR-Cas9 KO cell lines for different ceramide synthases stably expressing ER-SplitGFP.. Dot represents mean of 3 biological replicates. Mean ± SEM. **** p = < 0.0001 by One-way ANOVA and Tukey’s multiple comparison. **(D)** FACS assay measuring lysosomal transport of GM1-CTxB-pHrodo™ in A431 CRISPR-Cas9 KO cell lines for different ceramide. Dot represents mean of 3 biological replicates of 20,000 cells each. Mean ± SEM. **** p = < 0.0001 by One-way ANOVA and Tukey’s multiple comparison. **(E)** Lysosomal transport of GM1-CTxB-pHrodo™ in A431 WT cells mildly depleted of cholesterol as indicated. Dot represents mean of 3 biological replicates measuring 20,000 cells each. Mean ± SEM. **** p = < 0.0001 by One-way ANOVA and Tukey’s multiple comparison. **(F)** A431 WT cells were mildly depleted of cholesterol as indicated and retrograde transport was measured. Dot represents mean of 3 biological replicates measuring 20,000 cells each. Mean ± SEM. **** p = < 0.0001 by One-way ANOVA and Tukey’s multiple comparison.

To confirm these results, we took an orthogonal approach and generated A431 cells containing CRISPR-Cas9 deletion of CerS2, the ceramide synthase producing very long acyl chain ceramides (> C20), or deletion of CerS5, the synthase producing ceramides containing short acyl chain ceramides (≤ C16) (*61*). These deletions genetically bias the structures of the endogenous GM1 ceramide moieties in CerS2 and CerS5 KO cells as just described. The approach was validated by lipidomic profiling of sphingomyelin (SM). SM and GSLs share the same ceramide biosynthetic pathway and the abundance of SM in cell membranes allowed for the most conclusive analysis of endogenous ceramide structures. In wild type A431 cells, the major SM species contained C18:0 (45%), followed by C24:1 (15-18%), and C18:1 (10%) acyl chains (Suppl. Fig. 3C). Sphingomyelin containing ceramides with short acyl chains ≤ C16 (6-8%) were also reproducibly detected (n = 8 replicates). As expected in cells lacking CerS2, the SM species most strongly depleted were ceramides containing the very long acyl chains, but with concomitant increase in C18:0 and C18:1; while in cells lacking CerS5, the SM species most strongly depleted were the short acyl chains ≤ C16, and unexpectedly the C18 acyl chain species, with increases in C22:0 and C24:1.

When tested for GM1-dependent trafficking of CTxB, we found that the retrograde pathway was significantly reduced in those cells lacking CerS5 (Fig. 3C), consistent with 1) the loss of the short and C18:1 acyl chain ceramides that lack the C14*-motif; and 2) the increased expression of the very long chain saturated ceramide species C22:0 - including notably again those with an unsaturated bond positioned more deeply in the membrane (GM1 C24:1^Δ15^) - which contain the C14*-motif. We found a slight increase in retrograde trafficking for cells depleted in Cer2, consistent with depletion of only the very long chain ceramides and the modest increase in ceramides with C18:1^Δ9^ (Fig. 3C and Suppl. Fig. 3C). Again, control studies showed that endocytosis and EGF/LDL-R trafficking to the lysosome in CerS5 and CerS2 KO cells were not altered, or slightly enhanced (Fig. 2E, Suppl. Fig. 2G).

Opposite results were found for sorting CTxB to the acidic lysosome as predicted. This was measured by FACS using CTxB coupled to the pH-sensitive fluorophore pHrodo™. We found that CTxB-pHrodo™ fluorescence was reduced in the A431 cells deleted in CerS2, consistent with loss of very long acyl chain ceramides containing the C14*-motif and increase in the unsaturated C18:1^Δ9^ ceramide lacking the motif (Fig. 3D). For cells deleted in CerS5, which expressed higher levels of very long acyl chain ceramides (C20:0, C22:0, and C24:1^Δ15^), we found a slight increase in lysosomal transport - consistent with the enhanced expression of GM1 lipids containing the C14*-motif. Thus, the genetically modified cell lines phenocopied the trafficking of GM1 observed using exogenously applied and structurally defined GM1 species.

Cholesterol depletion had the same effect (Fig. 3E) – here we also observed an increase in retrograde trafficking to the ER (Fig. 3F). These results implicate a selective and cholesterol dependent process for retention within the body of the endosome followed by maturation to lysosomes for the GM1 ceramides that contain the C14*-motif (Fig. 3E and 3F). GM1 species lacking the C14*-motif escape this lysosomal sorting step and sort, perhaps passively by following bulk membrane flow, into endosome tubules serving the recycling and retrograde endocytic pathways.

### The C14* motif drives nanodomain assembly and segregation at the plasma membrane

The cholesterol dependency observed for C14*-motif based GM1 sorting to the lysosome is consistent with the idea that the mechanism of such sorting may involve assembly into membrane nanodomains. To test this, we imaged the GM1 C18:0 and C18:1^Δ9^ species in sorting endosomes artificially enlarged to enable visualization by over expression of dominant active Rab5Q79L (tagged with EGFP). We found clear separation of the two GM1 species in the plane of the endosomal membrane (Fig. 4A and B), and clear segregation between GM1 C18:0 (but not GM1 C18:1^Δ9^) and the Tfn-Tfn-R complex. Again, the unsaturated very long acyl chain GM1 C24:1^Δ15^, which also contains the C14*-motif, was found to behave like GM1 C18:0. These results imply nanodomain formation within the sorting endosome. Notably, the clear segregation of the two unsaturated GM1 species C24:1^Δ15^ and C18:1^Δ9^ into distinct endosomal nanodomains in live cells was not predicted by their phase behaviors in GMPVs (Suppl. Fig. 1B - D). In those studies, phase separation was dictated by the presence or absence of *cis*-double bond (kinked or straight shapes). In live cells, as evidenced here, we find the C14*-motif to be the decisive factor - not lipid shape.

**Fig. 4.**
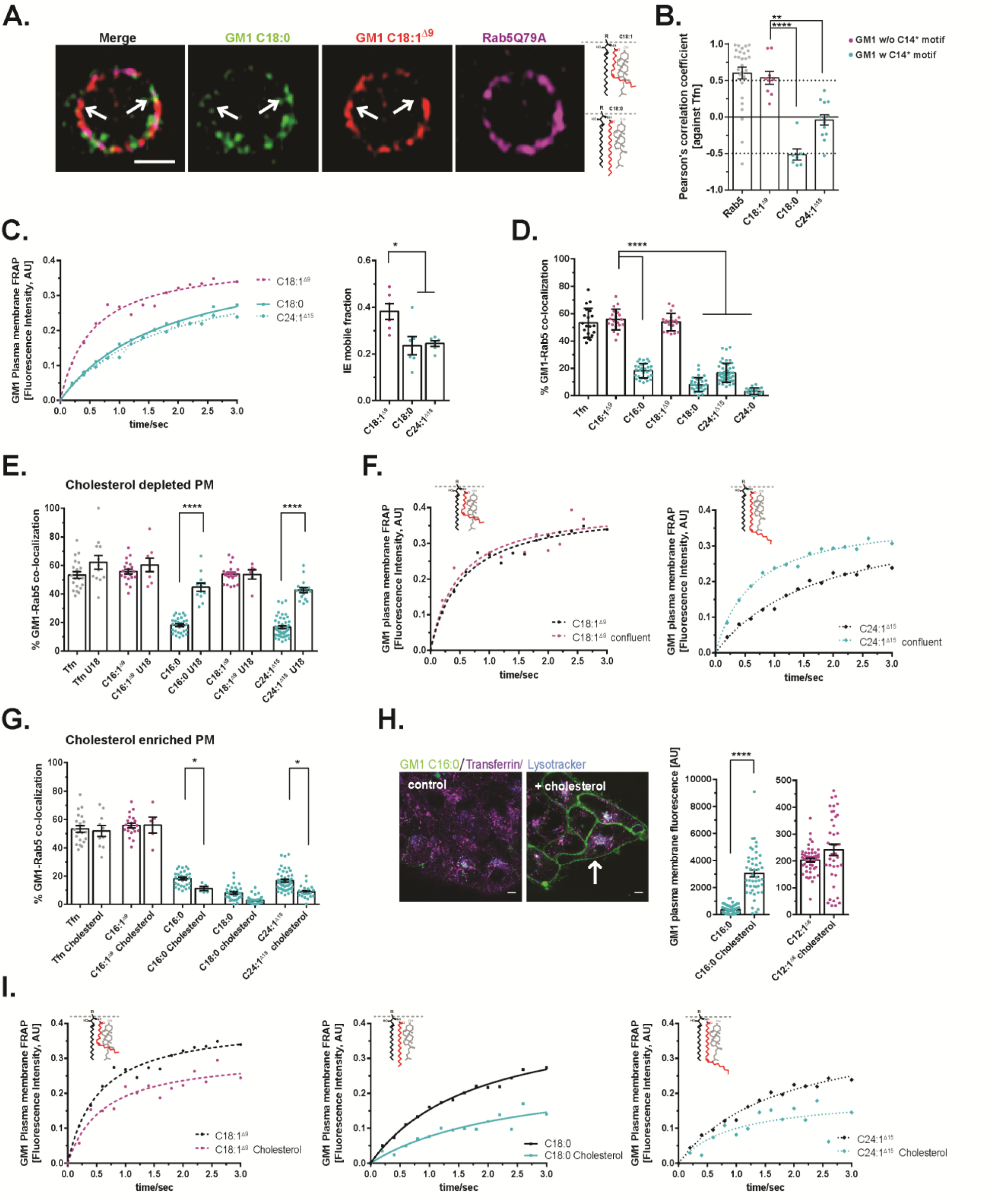
C14* motif drives GM1 segregation in endosomes and at the plasma membrane. **(A)** GM1 C18:0 (green) and GM1 C18:1^Δ9^ (red) were incorporated into A431 cells transfected with dominant active RAB5Q79L-GFP plasmid (magenta). Scale bar = 1 μm. **(B)** Pearson’s correlation coefficient against Tfn. Dot represents Pearson’s correlation coefficient for one enlarged RAB5 early endosome. GM1 species with C14* motif are displayed in turquoise and GM1 species without C14* motif in magenta. **** p = < 0.0001 by One-way ANOVA and Tukey’s multiple comparison.. **(C)** A431 WT cells were incorporated with the respective GM1 species and a plasma membrane ROI was photobleached immediately. Left panel: Fluorescence recovery after bleaching (FRAP) was recorded and mean grey values of one representative experiment are displayed. Right panel: Quantification of IE mobile fraction for FRAP experiments. Dot represents mean grey fluorescence for one FRAP experiment. Mean ± SEM. **** p = < 0.0001 by One-way ANOVA and Tukey’s multiple comparison. **(D)** Fluorescently labeled GM1 species were incorporated into genome edited SVGA cells expressing RAB5-GFP. GM1 - RAB5-GFP double positive vesicles were quantified. Mean ± SEM. **** p = < 0.0001 by One-way ANOVA and Tukey’s multiple comparison. **(E)** as in (D), but SVGA cells were mildly cholesterol depleted. **** p = < 0.0001 by One-way ANOVA and Tukey’s multiple comparison. **(F)** as in (C) but A431 cells were grown to over-confluency. **(G)** as in (D) but genome edited SVGA cells were incorporated exogenous cholesterol. Mean ± SEM. **** p = < 0.0001 by One-way ANOVA and Tukey’s multiple comparison. **(H)** Left panel: GM1 C16:0 (green) was incorporated into HT29 cells. GM1 trafficking was observed 30 min post loading in control cells or cholesterol treated cells. Scale bar = 1 μm. Right panel: Quantification of plasma membrane fluorescence of GM1 C16:0 and C12:1^Δ6^ incorporated into HT29 WT cells untreated or treated with exogenous cholesterol. Mean grey values of cell outlines for GM1 fluorescence were measured. Dot represents mean grey fluorescence for one cell outline. Mean ± SEM. **** p = < 0.0001 by One-way ANOVA and Tukey’s multiple comparison. **(I)** as in (C) but A431 cells were treated with exogenous cholesterol.

We were then curious to know if such segregation by nanodomain formation may also occur in the plasma membrane, before the lipids were internalized by endocytosis. To test this, we measured the diffusion rate of GM1 C18:0, C18:1^Δ9^, and C24:1^Δ15^ in plasma membranes of A431 cells using FRAP. We found slower rates of fluorescence recovery (FRAP) for the GM1 C18:0 and C24:1^Δ15^ species (which contain the C14*-motif) compared to GM1 C18:1^Δ9^ species (which lacks the C14*-motif) (Fig. 4C). Thus, even in the plasma membrane, the distinct behaviors among the different GM1 species appears to be dictated by the C14*-motif. The slower diffusion of the GM1 C18:0 and C24:1^Δ15^ species is consistent with their assembly in membrane nanodomains linked to the subcortical cytoskeleton (*62*).

To test if these differences in plasma membrane mobility correlated with differences in uptake, we measured the apparent rate of endocytosis. Here, we used SVGA cells endogenously expressing Rab5-GFP to measure the co-localization between the different GM1 species and Rab5 early endosomes. Uniformly, after short 5 to 10 min incubations, little co-localization with Rab5-endosomes was observed for GM1 ceramides containing fully saturated C16:0, C18:0, and C24:0 acyl chains and for the very long unsaturated acyl chain GM1 C24:1^Δ15^ (Fig. 4D) These ceramides all contain the C14*-motif. The result implicates slow rates of endocytosis, or very fast transition to Rab 7 late endosomes (or both) for these GM1 species. In contrast, we found a much higher degree of co-localization between the Rab5-endosome and GM1 ceramides containing the C16:1^Δ9^ and C18:1^Δ9^ acyl chains, which lack the C14*-motif - consistent with rapid rates of endocytosis, or efficient recycling (or both). The same results were obtained using Tfn, which we used as a control, phenocopying the C16:1^Δ9^ and C18:1^Δ9^ GM1 species (Fig. 4D).

When the SVGA cells were depleted of cholesterol, however, co-localizations of the GM1 C16:0 and very long C24:1^Δ15^ acyl chain species with Rab5 endosomes were increased (Fig. 4E). These lipids also displayed an increase in their PM mobility after cholesterol depletion as assessed by FRAP (Fig 4F, compare black with magenta lines and Suppl. Fig. 4A for quantification). When the SVGA cells were supplemented with additional exogenous cholesterol, however, the degree of colocalization with Rab5 endosomes was diminished for the same GM1 C16:0 and C18:0 species, and for the very long unsaturated acyl chain C24:1^Δ15^ species - compared to control untreated cells (Fig. 4G and Suppl. Fig. 4B). This phenotype induced by cholesterol addition was more readily apparent in HT29 cells, where PM localization was prominently marked at cell-cell contact sites. Here, these lipids appeared to be visibly “trapped” on the PM after cholesterol addition (Fig. 4G and H, and Suppl. Fig. 4D). Notably, the addition of cholesterol to SVGA cells had no detectable effect on the apparent efficiency of co-localization with Rab5-endosomes for the unsaturated GM1 species C16:1^Δ9^ and C18:1^Δ9^ (all lacking the C14*-motif) or for Tfn used as control. And we found no confounding effect of cholesterol treatment on fluid-phase dextran endocytic uptake or EGF-R/LDL-R sorting to the lysosome (Suppl. Fig. 4F).

Further evidence that excess membrane cholesterol caused retention of GM1 species containing the C14*-motif on the PM was obtained in A431 cells as assessed by FACS (Suppl. Fig. 4E) and by confocal microscopy (Fig. 4H). - and with no evidence for co-localization within Tfn-positive endosome recycling tubules (Suppl. Fig. 4D). We also found a striking reduction in PM diffusion rates for the C18:0 and C24:1^Δ15^ species in A431 cells treated with exogenous cholesterol - as assessed by FRAP (Fig. 4I and Suppl. Fig. 4A and see (*63*)). These results suggest nanodomain assembly for the C16:0, C18:0, and notably C24:1^Δ15^ GM1 lipids.

Thus overall, we found that the different GM1 species are sorted within live cells based on the C14*-motif. These GM1 species were retained within the body of the sorting endosome for maturation along the lysosomal pathway. And they exhibited dependence on membrane cholesterol for their sorting and biophysical behaviors, consistent with an affinity for assembly into membrane nanodomains (Fig. 5). In contrast, the GM1 species containing mono-unsaturated acyl chains of 12 to 22 hydrocarbons atoms in length (thus lacking the C14*-motif) did not display dependence on cholesterol for their membrane behaviors. And unlike the fully saturated GM1 species, these GSLs entered endosome sorting tubules of the recycling and retrograde endosome pathways.

**Fig. 5.**
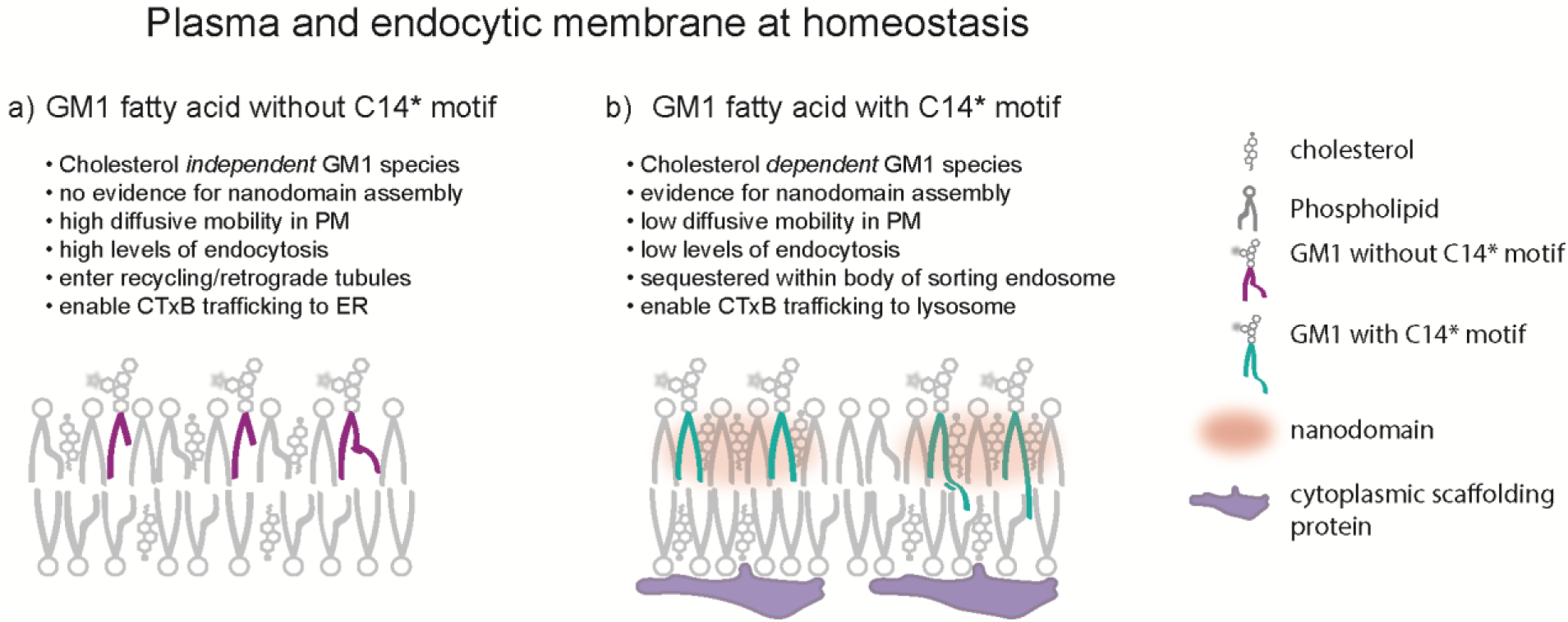
Model for cholesterol-ceramide structure based GM1 sorting. a) GM1 species with short or unsaturated acyl chains (≤ C14:0 or C22:1^Δ13^) lacking the C14* motif (magenta) are unable to assemble with cholesterol into membrane nanodomains. Their membrane and endocytic trafficking phenotypes are therefore independent of membrane cholesterol. b) GM1 species with long or very long unsaturated acyl chains (≥ C16:0 and C24:1^Δ15^) containing the C14* motif (turquoise) allow for close packing with cholesterol and therefore assembly into membrane nanodomains (or rafts).

## Discussion

The results of our studies show that mammalian cells sort GM1 sphingolipids between endosomes and endosome sorting tubules, and within the plane of the PM, based on their ceramide acyl chain structure. This is explained by differential association with membrane cholesterol - likely mediated by lipid nanodomain formation (Fig. 2C and schematic in Fig.5). Lipid shape on its own (*55, 64–66*) cannot fully explain the selective entry of the different GM1 species into the narrow highly curved endosome sorting tubules, or their retention in the body of the sorting endosome. The differential sorting phenotypes observed were strictly dependent on membrane cholesterol content and ceramide structure, and the same structure-function relationships operated across multiple cell types known to vary in their overall membrane thickness, fluidity, and cholesterol content (our unpublished data and (*67–69*)). These results define a structural organizing principle for complex sphingolipid behavior in native heterogeneous and dynamic live cell membranes. The complex sphingolipids critically require more than 14 saturated hydrocarbons in the ceramide acyl chain extending from the C1-amide at the water-bilayer interface for optimal association with cholesterol. And this decisively affects their direction of endosome sorting. GSL species lacking these features escape the apparently active lysosomal sorting step and enter endosome sorting tubules of the recycling and retrograde pathways instead. These observations experimentally verify in live cells one of the underlying and fundamental biophysical processes hypothesized to shape lipid trafficking, membrane architecture, and cell function. It is possible the C14*-motif may also explain the differential sorting of ceramide-linked proteins in the secretory pathway recently observed in yeast (*41*).

In nature, the *cis*-double bond in the GM1 species containing mono-unsaturated acyl chains of 16 to 22 hydrocarbons atoms in length is positioned at carbon atom 9 or 13 (Δ^9^ and Δ^13^) extending from the amide linkage to sphingosine, and thus located within the outer leaflet of the membrane bilayer. Within this area, the “kink” in the acyl chain caused by unsaturation interferes with alignment against the flat and rigid sterol structure of cholesterol (*43, 70–74*). These GM1 species entered endosome sorting tubules of the recycling and retrograde endosome pathways, and their trafficking was not dependent of cholesterol. On the other hand, the GM1 species containing the very long mono-unsaturated acyl chains (C24:1^Δ15^ and C26:1^Δ17^) trafficked to the lysosome - phenocopying the fully saturated GM1 species even though they also contained a *cis*-double bond “kink” in their ceramide. Here, the structural analysis was most informative. The *cis*-double bond in these native acyl chains is located at Δ^15^ or Δ^17^, below the stretch of 14 saturated hydrocarbons from the amide linkage at the membrane surface (*44, 70*), and below the expected interface with the sterol rings of cholesterol (*43*). Thus, the acyl chains of these GM1 species recapitulated exactly the minimal saturated hydrocarbon chain length that we found required for endosome sorting of the GSLs to the lysosome. Notably, the membrane behavior of these very long unsaturated GM1 species in live cells was strictly dependent on membrane cholesterol, analogous to the fully saturated acyl chain GM1 species. Two species GM1 C14:0 and GM1 C22:1^Δ13^ had intermediate phenotypes for cholesterol dependence - these ceramides have acyl chain lengths that border on the minimal C14*-motif required for functional association with cholesterol.

The strong dependence on cholesterol for the membrane behaviors of the saturated and very long chain unsaturated GM1 species conforms to central tenants of the lipid raft hypothesis. The raft hypothesis posits that certain lipids and cholesterol phase separate into ordered and tightly packed domains, mediated through hydrogen bonding and van der Waals forces, and shaped by cellular protein scaffolds, including by bonding to the cortical actin cytoskeleton (*18, 20, 75*). For the saturated and very long chain unsaturated GSLs, we find evidence for their assembly into membrane nanodomains: perhaps best evidenced by the segregation of these GM1 species in separate regions of artificially enlarged early endosomes. Our results are consistent with another structural factor implicated in nanodomain assembly for GPI-anchored proteins and recently for GM1 ((*76*) and our unpublished results, Arumugam et al.): the need for direct extension of outer membrane lipids into the inner-membrane lipid leaflet for assembly with phosphatidyserine and the cortical actin cytoskeleton. In those studies, we found that the ceramide C14*-motif was required for features of nanodomains with linkage to the cortical cytoskeleton as assessed by fluorescence correlation microscopy. Nanodomain formation, and GM1 trafficking, may also be affected by extracellular factors that induce lipid clustering (*14*), as we found for CTxB (*40*).

The structural diversity of ceramide moieties comprising the complex sphingolipids that occur in nature is perhaps best defined for sphingomyelin, which is the most highly prevalent complex sphingolipid, and thus most easily and reproducibly studied. In our hands, and in the cell-type examined, we found that the predominant ceramide moieties in sphingomyelin contained C18:0, C18:1^Δ9^, C24:0, and C24:1^Δ15^ acyl chains. This, and published lipidomic analyses of sphingomyelin in other cell types, suggest the evolution of two ‘bi-modal’ structure-function distributions for the complex sphingolipids: one between long and very long ceramide acyl chains as proposed before [(*46*)61-64], and the other between saturation and unsaturation as we find here. Experimentally, these two bi-modal distributions were evident in our structure-function studies on GM1. In the case of unsaturation, the different membrane behaviors were most readily apparent for the saturated GM1 C18:0 and the unsaturated C18:1 species. But this dichotomy of saturation and unsaturation was not absolute - it also depended on acyl chain length. In the case of GM1 species with very long C24:0, and C24:1^Δ15^ acyl chains, for example, these GSLs behaved mostly in the same way - even though one was saturated and the other was not. Thus, acyl chain length in the native complex sphingolipids appears to over-ride the functional impact of unsaturation, as long as the *cis*-double bond is positioned below a stretch of saturated hydrocarbons of sufficient length (beyond the C14*-motif). The long and very long acyl chains may also enable interdigitation with the contralateral lipid leaflet (*77–79*). Lipid shape, on its own, does not appear to act decisively for entry of the GSLs into endosome sorting tubules, or for their endosomal sorting – contrary to the impact of acyl chain unsaturation on L_d_ and L_o_ phase separation and nanodomain assembly in GPMVs.

We also note that some very long unsaturated acyl chain GM1 species, typified by GM1 C24:1^Δ15^, exhibited graded membrane behaviors that bridged the dichotomy between saturation and unsaturation seen in the GM1 C18:0 and C18:1 pair. These very long acyl chain GSLs sorted predominantly to the lysosome, but a significant portion sorted also into the recycling pathway - in contrast to their fully saturated counterparts. In addition, GM1 C24:1^Δ15^ had a restricted but still higher degree of apparent endocytic uptake compared to its fully saturated counterpart. Thus, the very long chain unsaturated GSLs appear to occupy a functionally intermediate space. They may have been evolutionarily conserved according to their graded membrane behaviors in membrane sorting so as to more readily enable differential responses to small alterations in membrane structure and composition caused by cell stress, cell differentiation, or even to the normal dynamic changes in membrane content produced by trafficking from one organelle to another in cells at steady state (*80*).

In summary, our studies show that the membrane behaviors of the complex sphingolipids are defined by two general rules of ceramide structure - hydrocarbon acyl chain length and unsaturation. This includes the position of the *cis*-double bond with respect to the outer leaflet of the membrane lipid bilayer and how acyl chain length and saturation affect the ability of GSLs to associate with cholesterol and possibly to form nanodomains. These basic structural principles are of general importance to cell and tissue biology, as the segregation and behavior of lipids has fundamental physiological consequences for membrane organization and the plasticity required to accommodate the diversity of membrane dynamics underlying cell functions.

## Material and Methods

Dimethylformamide (DMF), dimethylsulfoxide (DMSO), methyl-β-cyclodextrin (MβCD), soluble cholesterol complexed to MβCD, defatted bovine serum albumin (dfBSA), fetal bovine serum (FBS), fetal bovine serum depleted of HDL (HDL-), Simvastatin (statin), T0901317 (T09), U18666A (U18) and filipin III (filipin) were purchased from Sigma Aldrich (St. Louis, MO).

Dulbecco’s modified Eagle’s medium (DMEM) with or without phenol red, OptiMEM®, Lipofectamine® 2000, Laurdan (6-Dodecanoyl-2-Dimethylaminophthalene), *FAST*Dil™ Dil^Δ9,12^- C_18_(3), CIO_4_ (1,1’-Dilinoleyl-3,3,3’,3’-Tetramethylindocarbocyanine Perchlorate), *FAST*Dio™ DiO^Δ9,12^-C_18_(3), CIO_4_ (3,3’ Dilinoleyloxacarbocyanine Perchlorate)), various AlexaFluor™ (- 405, - 488, - 568, or - 647 nm) or pHrodo™Fluor-NHS esters and –azide dyes, AlexaFluor™ and pHrodo™Fluor conjugates, Dextran Pacific Blue™ (3000 MW) and LysoTracker™ were purchased from Thermo Fisher Scientific (Waltham, MA).

### Cell culture

Human epithelial A431, HT29, CaCo and Hela cells, as well as HEK293T cells were originally purchased from the American Tissue Culture Collection (ATCC, VA). Cells were maintained in Dulbecco’s modified Eagle’s medium (DMEM) containing 10% FBS with penicillin and streptomycin. Cultures were split two days prior to experiment to achieve the desired confluency state. The CRISPR-Cas9 SVGA Rab5-GFP cell line was maintained in MEM containing 10% FBS.

For cholesterol addition and/or depletion assays, cells were cultured for 24 h in [50 – 100 nM] T0901317, or for 12 h in [1 −2 μg/ml] U18666A in full medium and for 24 h for [10 μM] simvastatin in serum depleted of HDL. Soluble cholesterol complexed to MβCD [120 μg/ml] was given in full media for 30 min or the indicated time. Efficacy of treatment was determined by fixing treated cells in 4 % paraformaldehyde for 1 h and subsequently staining cells with [0.05 mg/ml] filipin III to determine plasma membrane cholesterol contents by fluorescence. Plasma membrane fluorescence intensity was measured using the Fiji (ImageJ, National Institutes of Health, Bethesda, MD, USA) segmented line tool to outline plasma membrane and mean grey values were obtained.

To obtain enlarged Rab5 early endosomes, A431 cells were transfected using Lipofectamine**®** 2000 and OptiMEM**®** transfection method according to manufacturer’s instructions and 500 – 1000 ng of the Addgene plasmids (#28046, #35138).

### Generation of CRISPR-Cas9 KO cell lines

CRISPR-Cas9 KO cell lines were generated using Addgene plasmid lentiCas9-Blast (#52962) for stable integration of Cas9 endonuclease and guide sequences were cloned into Addgene plasmid lentiGuide-Puro (#52963) against ceramide synthase 1, 2 and 5 (CerS1, 2 and 5), UDP-glucosyl-ceramide glucosyltransferase (UGCG), and serine palmityl transferase 2 (SPTLC-2) (see Table 1). Plasmids were verified by sequencing using IDT technologies (Coralville, IA). [1.5 μg] plasmids were co-transfected with packaging plasmids pVSVg [0.5 μg] and psPAX2 [1 μg] Addgene plasmids #8454, #12260 in the presence of polyethylenimine (PEI) into HEK293T cells for lentivirus production. 48 h post transfection virus was harvested and mixed with respective cell line. Clones were selected for 7 days on [10 μg/ml] Blasticidin and/or [1 μg/ml] Puromycin. KO cell lines were tested for CTxB binding by FACS and low binding population was sorted.

**Table 1.**
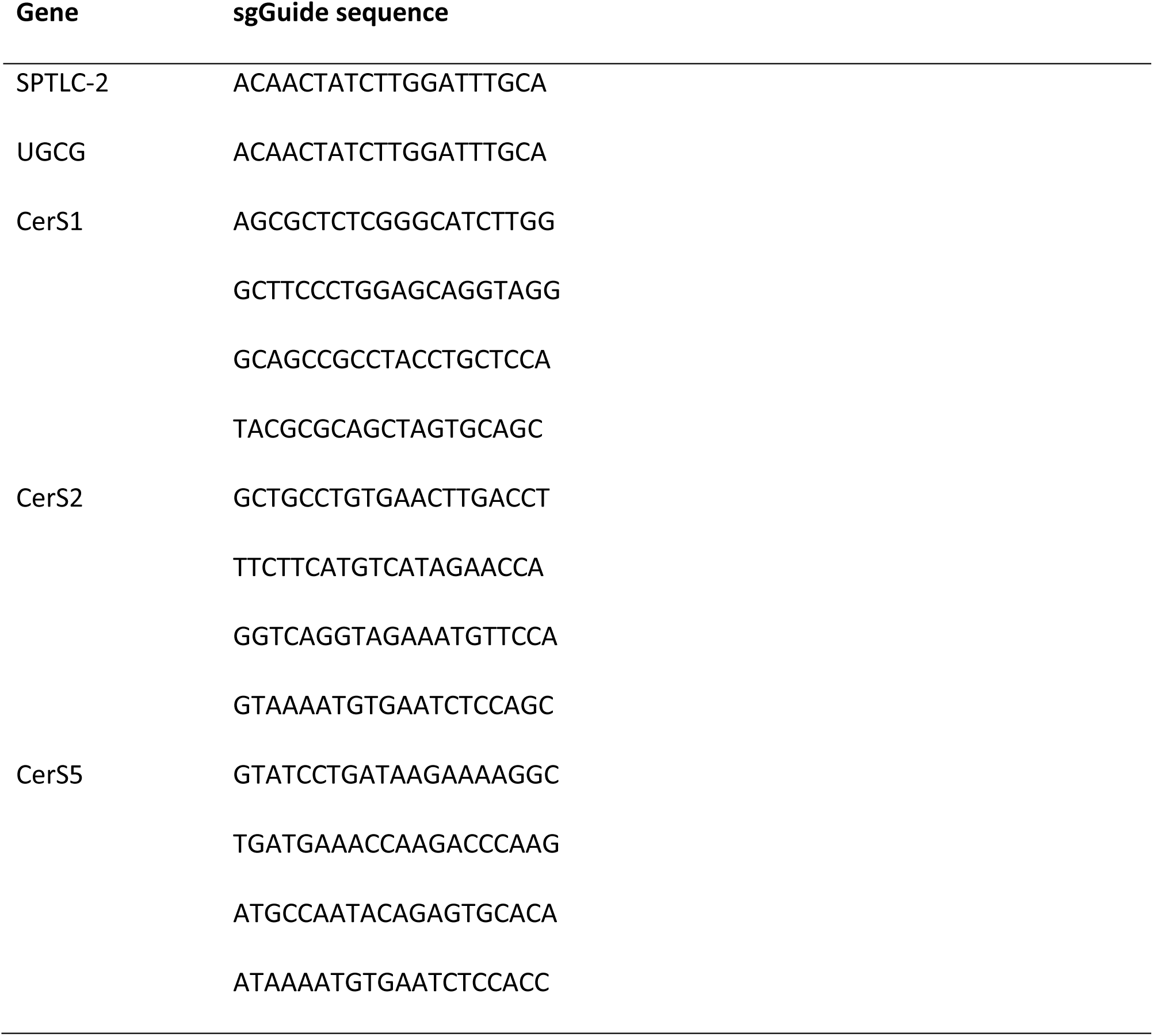
CRISPR KO guide sequences Guide sequences were used from AVANA sgGuide library (*85*).

Cell lines carrying an ER version of Split-GFP were constructed according to Luong et al. (*60*) and selected on Hygromycin [50 μg/ml] for 7 days.

### Synthesis of peptide and fluorophore-labeled GM1 species

Gangliosides of various fatty acid species were supplied by Prof. Sandro Sonnino (U. Milan, Italy). Reporter peptide containing modified functional residues were custom synthesized by New England Peptide (Gardner, MA) as all-D peptides with the following amino acid sequence: propargylglycine-k(ε-biotinoyl)(ds)g(dy)g(dr)g(ds)g-(kaoa)-amine. Synthesis of peptide-lipid and fluorophore labeled conjugates was accomplished with minor modifications according to Garcia-Castillo et al., te Welscher *et al.* and Chinnapen et al. (*40, 47, 48*).

In a typical 2 ml reaction, 2 mg (approximately 1300 nmoles) of ganglioside was oxidized with sodium periodate (13 mmoles) in oxidation buffer (100 mM sodium acetate pH 5.5, 150 mM NaCl) for 30 min on ice. The reaction was quenched by addition of glycerol (5% final) and desalted using a Bond Elut SepPak C18 cartridge (Agilent, MA). Methanol was used to elute oxidized GM1 species from the column and was removed by Speed Vac concentration (Savant). The oxidized product was then reconstituted in 2 ml PBS pH 6.9 in the presence of 10% DMF and reacted with 2700 nmoles of aminooxy-containing peptide in the presence of 10 mM aniline (Dirksen and Dawson, 2008). The reaction was incubated for 20 h at room temperature with mixing on a nutator. The precipitate was separated from the solution by centrifugation, and then resuspended in 400 μl 50% isopropanol/water after brief sonication. PBS pH 6.9 was added (200 μL) along with 4.8 mmoles of sodium cyanoborohydride and incubated for 3 h to reduce the oxime bond. Lipid-peptide conjugates were purified by semi-preparative HPLC, and confirmed by either MALDI-TOF (AB Voyager), or ESI LC-MS (Agilent, MA).

Peptides were labeled with Alexa fluorophore via copper mediated Click chemistry. 320 mM peptide-lipid fusions containing an N-terminal alkyne residue (propargylglycine) were reacted with equimolar concentrations of AlexaFluor™-azide under the following conditions: 50 mM Tris-Cl, 5 mM copper (II) sulfate, 100 mM sodium ascorbate, 37 mM (Tris[(1-benzyl-1H-1,2,3- triazol-4-yl)methyl]amine, TBTA in DMSO/t-butanol 1:4) 1 mM (Tris(2-carboxyethyl) phosphine hydrochloride, TCEP – Sigma) for 16 h at room temperature on a nutator. Products were purified by HPLC and confirmed by mass spectrometry. Products were lyophilized and stored at - 20°C or - 80°C for long term storage.

Working stock solutions were prepared in 33% DMF/ddH20

Alternatively, fatty acyl variant gangliosides were fluorophore labeled directly via the oligosaccharide head group following the oxidation of the ganglioside. The oxidized gangliosides were resuspended in phosphate buffered saline pH 7.4 (PBS) and reacted with 1 mole AlexaFluor™-azide (Invitrogen, OR) for 16 h. Ten moles sodium cyanoborohydride was added to the reaction for 15 minutes and the mixture was desalted on Bond-Elute cartridges. Ganglioside-fluorophore conjugates resulted in a different mobility band visible by TLC and were purified using preparative-TLC plates. Extraction from silica scrapings was performed, and the conjugates further purified by reversed phase C18 cartridges.

### FACS based assay for lipid recycling, retrograde and lysosomal trafficking

Cell lines were plated on 96 well plates and grown to 70% confluency (unless otherwise stated) with or without treatment. Cells were washed and equilibrated with serum-free DMEM (no phenol red) for 5 min at 37°C. GM1 species were diluted to four different concentrations of the respective peptide-labeled GM1 species in serum-free DMEM with equimolar ratio of dfBSA (1:1 lipid:dfBSA). Lipids were loaded for 10 min at 37°C and washed of in serum-free DMEM (no phenol red). To measure the amount of ‘loaded’ GM1, cells were trypsinized for 5 min at 37°C, then chilled to 4°C and stained with respective streptavidin-AlexaFluor™ (1:2500, 2 mg/ml) in PBS containing 5% BSA for 15 min/4°C. Unbound streptavidin was washed off twice with PBS containing 5% FBS and cells were FACS analyzed immediately, using a FACS Canto II (BD, Biosciences, NJ). Lipid+ gate was determined against streptavidin stained control cells, not loaded with GM1. The percentage of positive cells and their median fluorescence were recorded. To compare the recycling efficiency between the different GM1 variants, the respective concentrations resulting in equal median fluorescence or equal % of cells for the ‘load’ were chosen. To obtain ‘chase’ samples, the lipid was washed off and cells were subsequently incubated for 3 h in DMEM containing 5% FBS in the presence of Dextran-Fluor (500 μg/ml) and transferrin-AlexaFluor™ (1 μg/ml). Cells were then washed and trypsinized and stained with streptavidin-AlexaFluor™ as described above. To compare recycling rates between different treatments, recycling rates (median fluorescence) for non-treated or WT cells were normalized to 1 and fold change difference to treated samples calculated.

To determine retrograde trafficking of CTxB-SplitGFP, native (non-peptide labeled) GM1 species were loaded as above into either Hela cells or an A431 CRISPR-Cas9 UGCG-KO cell line. ‘Load’ amount of lipid was determined using 5nM fluorescently labeled CTxB instead of streptavidin as described above. ‘Chase’ samples were incubated in the presence of 10 nM CTxB-SplitGFP for the indicated amount of time in DMEM containing 5% FBS. For retrograde trafficking in CRISPR-Cas9 KO cell lines, cell surface GM1 amount was determined using 5nM fluorescently labeled CTxB as described above and CTxB-SplitGFP values were normalized to cell surface GM1.

To measure lysosomal trafficking of GM1-CTxB complex, pH sensitive CTxB-pHrodo™ was used. Cells were chilled to 4°C and stained using 5 nM CTxB-pHrodo™ in PBS containing 5% BSA. Unbound CTxB was washed off and cells were incubated for the indicated amount of time at 37°C in in DMEM containing 5% FBS. Cell surface GM1 was determined using 5nM fluorescently labeled CTxB as described above and CTxB-pHrodo™ values were normalized to cell surface GM1.

Two biological replicates per assay were performed and a total of N = 10,000-20,000 single cells recorded per treatment for all FACS based assays.

### Lysosomal trafficking of EGF receptor and LDL receptor

Cells were grown as described above. To determine the lysosomal fraction of different membrane protein receptors, cells were loaded with their respective ligands at (2 - 5 μg/ml), labeled either as pH sensitive (phrodo™, FITC) or pH resistant (AlexaFluor™/Bodipy for LDL). ‘Load’ was determined by incubating pre-chilled cells for 10 min at 4°C with pH resistant ligand. Cells were subsequent washed in PBS and detached using Cell Dissociation Buffer (Gibco). Lysosomal fraction was determined by loading cells with pH sensitive ligand as above; after washing off unbound ligand, cells were moved to 37°C for 2 h in the presence of fluorescent dextran to normalize endocytosis rates. Cells were trypsinized and subsequently analyzed by FACS.

### Lipid loading, imaging and quantification for Airyscan or lattice lightsheet imaging

Cell lines were plated on either 35 mm MatTek petri dishes (Ashland, MA) containing glass bottom for Airyscan confocal microscopy or on 3 mm cover slips for Lattice Lightsheet microscopy and grown to 70 % confluency with or without treatment. Fluorescent GM1 variants were added as described for the FACS based assays for lipid recycling unless otherwise noted. To determine subcellular location, lipids were allowed to traffic for the indicated amount of time at 37°C in the presence of fluorescent transferrin (1 μg/ml) and lysoTracker™ (2 nM) or dextran (1 mg/ml) to demarcate the respective organelles. Sorting endosomes were defined as vesicles containing lipid as well as transferrin and dextran. Sorting endosomal tubules were defined by presence of transferrin emerging from triple positive vesicles.

For lactose and sucrose addition experiments A431 cells were incubated for 30 min in the presence of 100 mM lactose or sucrose prior/during and after GM1 loading.

Airyscan imaging was performed using Zeiss LSM 880 microscope with Airyscan detector and Plan-Apochromat 63x oil immersion objective with NA = 1.4. Airyscan processing was performed using the Zeiss ZenBlack software.

Quantification of GM1 in recycling endosomal tubules:

Region of interest (ROI) was determined, containing approximately 3 - 6 triple positive vesicles and the ROI was thresholded using image quantification software ImageJ (Otsu). GM1 presence/absence (1/0) within transferrin positive tubules was recorded and % for the ROI calculated.

Tubule over vesicle fluorescence quantification:

For GM1 C14:0 and C22:1^Δ13^ as well as for samples treated with cholesterol lowering molecules, or cholesterol addition, fluorescence grey values for the vesicle and tubule were recorded, background fluorescence was subtracted and a ratio was determined by dividing the mean grey value obtained for the tubule by the mean grey value obtained for the vesicle (Fig. 1H). The same analysis was performed for Fig. 2G and Suppl. Fig. 4D.

Quantification of endocytic rates:

Endocytic rates were assayed using CRISPR-Cas9 modified SVGA cell line containing an endogenous, RAB5-EGFP variant (*81*). GM1 variants were loaded as described above, washed off and immediately transferred to the microscope. Cells were imaged for approx. 5 min each.

Vesicles were counted using ImageJ software and quantification was based on at least 5 biological replicates.

GM1 - Tfn co-localization in Rab5 enlarged early endosomes:

Using Fiji segmented line tool Rab5 early endosomes were traced using Rab5 fluorescence as marker and mean grey fluorescence intensity profiles of GM1 variants and Tfn were exported into GraphPad Prism. Pearson’s correlation coefficient was calculated between different GM1 or Rab5 fluorescence profiles against Tfn fluorescence profile.

### Fluorescence recovery after bleaching (FRAP)

Cell lines were plated on 35 mm MatTek petri dishes containing glass bottom 48 h prior to experiments and treated as described above. GM1 lipids were incorporated into plasma membranes as described above.

Cells were immediately transferred into preheated chamber for bleaching using the following settings: 80 – 100 % laser intensity, pixel dwell time of approx. 170 μsec, membrane fluorescence was bleached to about 20 - 40% of the original fluorescence intensity. Three ROIs of similar size were chosen, one background ROI (ROI_bg_) and two ROIs within the PM of a given cells, with one of the latter being bleached (ROI_b_), while the other served as reference photobleaching control (ROI_nb_). To establish the baseline, three – five frames were collected before the bleaching event and to capture the recovery, at least fifteen frames were acquired every 150 - 300 msec for a total time of approx. 10 sec post bleaching event.

FRAP curves were determined using the following formulas with pb = pre bleached fluorescence (*82*):

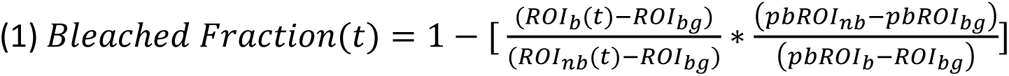

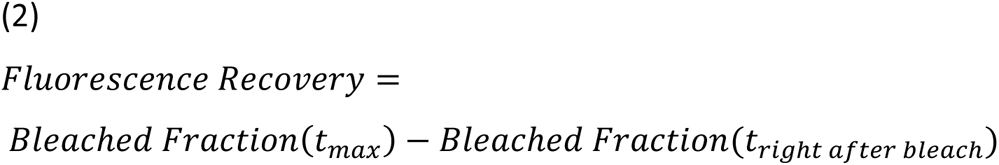

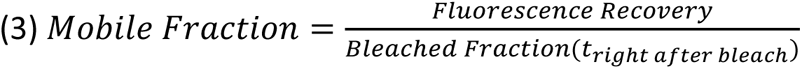

### GPMV generation

Cells were seeded two days prior to assay on 12 well plates to reach desired confluency. Native or fluorescently labeled GM1 species were added as described above, and a phase marker (*FAST*DiO™ and *FAST*DiL™, 0.5 mg/ml − 1. 1:1000 in PBS) was added for 10 min. Phase marker was washed off three times using PBS, then cells were equilibrated by washing twice in GPMV buffer (50 mM Tris, pH = 8, 150 mM NaCl, 50 mM CaCl_2_). Vesiculation was achieved according to (*50*) by using 2 mM DTT and 0.07 % PFA in GPMV buffer and cells were incubated for 1-3 h at 37°C (depending on the cell type). Supernatant containing GPMV vesicles were harvested and sedimented 5 min/ 500 g. Vesicles were chilled to 10°C and imaged. L_d_ phase was determined by the phase marker and L_o_ partitioning coefficient (ordered phase partitioning coefficient = I_lo_) was obtained using the following formula: K_raft_ = I*_Lo_*/I*_Ld_*. A K*_lo_* > 1 indicates ordered phase preference.

For membrane fluidity analysis, GPMVs were harvested from respective cells and/or treatments as above and subsequently stained using [0.5 μM] laurdan for 3 h at RT. Dye was washed off twice and vesicles were chilled to 10°C prior to imaging. Imaging was achieved by using spectral imaging with confocal microscope Zeiss LSM 880 using Plan-Apochromat 63x oil immersion objective (NA 1.4). C-laurdan was excited at 405 nm and the emission was collected with a multichannel spectral detector (PMT) in the range 430 - 450 nm and 480 - 500 nm simultaneously. The polarity index, generalized polarization (GP) was calculated according to Steinkuehler *et al.* (*68*) using the following formula: GP = (I_440_ − I_490_)/(I_440_ + I_490_) where I_440_ and I_490_ are the fluorescence intensities at 440 nm and 490 nm, respectively.

### Lipid extraction and Lipidomic profiling of A431 WT and CRISPR-Cas9 KO cell lines

500,000 cells per biological/technical replicate were washed twice with PBS and then 0.25 x PBS to remove media components and transferred into glass vials. Cells were then pelleted to remove remaining supernatant and flash frozen in liquid nitrogen and pellet was subsequently lyophilized. Lipid extraction and lipidomic profiling was performed according to Liaw et al. and Kumar et al. (*83, 84*). In brief, the lyophilized pellet was homogenized in 1 ml ddH_2_O, subsequently 2.2 ml methanol and 0.9 ml dichlormethane was added. Suspension was incubated o/N at −20C, before 20 μl Avanti Splash mix was added, mixture was vortexed and incubated for 10 min at RT. Then 1 ml ddH_2_O, 0.9 ml dichloromethane and 0.5 ml methanol were added. Solution was centrifuged at 1200 rpm for 10 min and lower phase collected into fresh glass tube. 2 ml dichloromethane were added to the aqueous phase to extract any remaining lipids, mixture was centrifuged as above and the organic phase was pooled with the first one.

Solvent was evaporated in Genevac.

Unbiased MS/MSALL shotgun lipidomic profiling was performed on Sciex 5600 TripleTOF mass spectrometer.

### Software and Statistical analysis

Imaging data was analyzed using either Zeiss ZENblack, Fiji or Aiviva 9 (Drvision Technologies, Bellevue, WA) software. Lattice lightsheet data obtained at the Janelia Farms Advanced Imaging center (AIC), deskewed and decovoluted using customized Matlab™ software.

Schematic Model in Fig. 1A was drawn using BioRender.

Mean grey fluorescent values of microscopy experiments or mean fluorescence data obtained from FACS experiments was transferred to GraphPad Prism software (San Diego, CA) for graphing and statistical analysis. Statistically significant difference between treatments was tested by One-way ANOVA with Tukey’s multiple comparison; alternatively unpaired t-test was applied when indicated with n.s. p > 0.05, *p < 0.05, **p < 0.01 ***p < 0.001 and ****p < 0.0001.

## Acknowledgements

We thank the Janelia Farms Advanced Imaging Center for providing access and support to their Lattice Lightsheet microscope. Imaging data used in this publication was produced in collaboration with the Advanced Imaging Center, a facility jointly supported by the Gordon and Betty Moore Foundation and HHMI at HHMI’s Janelia Research Campus. We further thank the Harvard Center for Biological Imaging, Boston Children’s Hospital Cell Function and Imaging Core and HDDC Imaging Core for infrastructure and support.

We thank Prof. Dr. Kirchhausen (Boston Children’s Hospital, Harvard Medical School, USA) for providing the SVGA-Rab5-GFP cell line, as well as Prof. Dr. Waldor and Dr. Alline Pacheco (Brigham and Woman’s Hospital, Harvard Medical School, USA) for providing one CRISPR-Cas9 UGCG and SPTLC-2 KO plasmids and HT29 Cas9 positive cell line.

We further thank Dr. Daniel Chinnapen, Dr. Richard Duclos, Dr. Phi Luong and Dr. Jamie LeBarron for helpful discussions during the preparation of this manuscript and Profs. Thiagarajah (Boston Children’s Hospital, Harvard Medical School, USA), Johannes (Institut Curie, France) and Kenworthy (University of Virginia, School of Medicine, USA) for critically reading the manuscript.

This work was supported by grants R37 DK048106 and RO1 DK104868 to WIL, and P30 DK034854 to the Harvard Digestive Disease Center

## Author contributions

S.S.S and W.I.L. conceive study; S.S.S, R.T., W.I.L. designed experiments and resources; S.S.S, R.T. and K.K. performed experiments; S.S.S, R.T., and M.A. analyzed data. S.S.S. and W.I.L., wrote manuscript.

## Supplementary Figures

**Suppl. Fig. 1.**
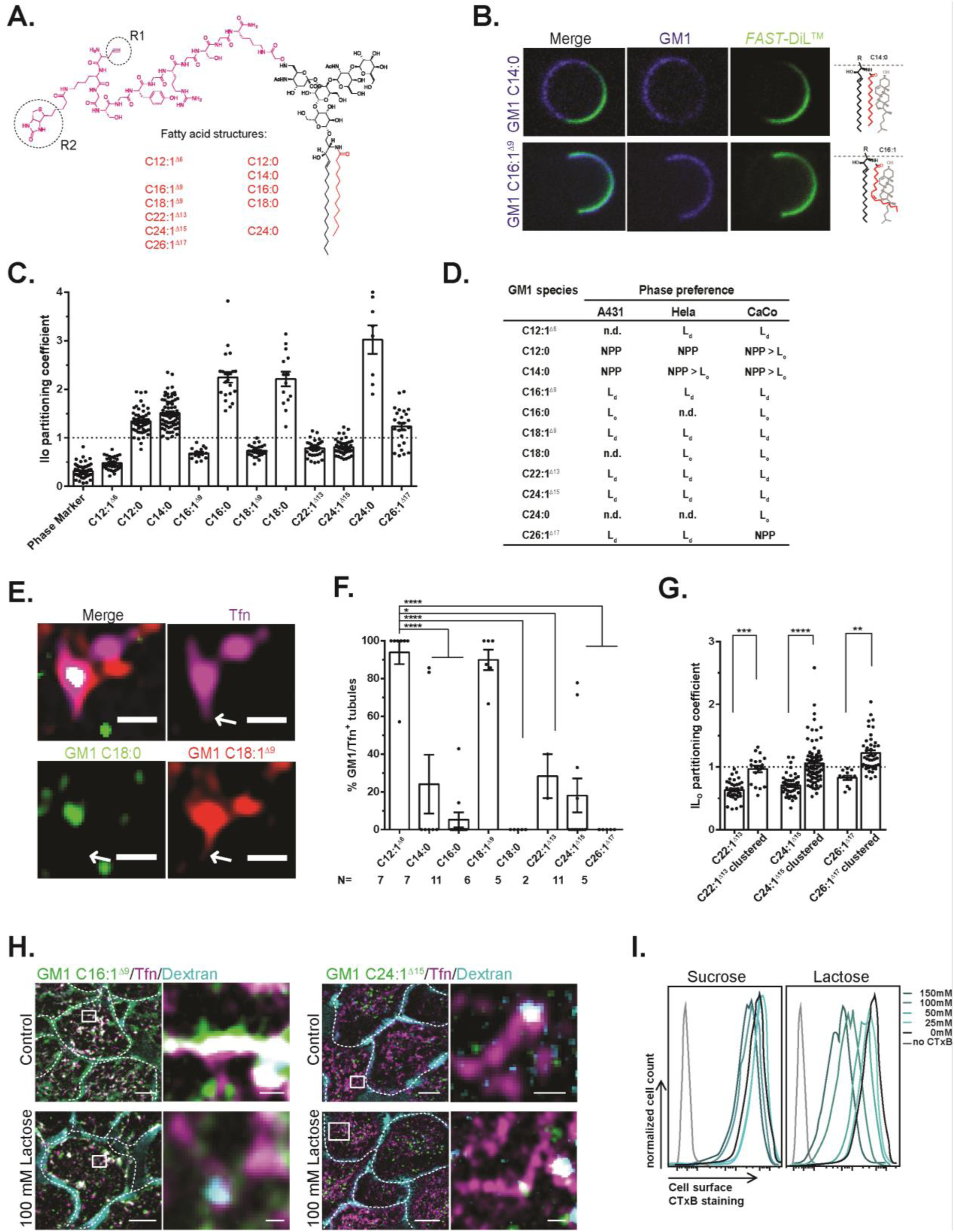
Ceramide structure specific biophysical behavior and cellular sorting of GM1. **(A)** Structure of the linker ‘reporter’ peptide (magenta) coupled to the sialic acid of GM1 oligosaccharide (black) with functional biotin (R2) for streptavidin-AlexaFluor™ detection and alkyne group (R1) for direct AlexaFluor™ azide conjugation. Sialic acid is also used for direct AlexaFluor™ - GM1 conjugates. Different fatty acyl chain structures used in the study are displayed in red. **(B)** GPMV from CaCo BBE WT cells loaded with fluorescent GM1 C14:0 and C16:1^Δ9^ and *FAST*-DiL™ phase marker. Exemplary images shown. **(C)** Phase preference (IL_o_ partitioning coefficient) for the library of GM1 species. Dot represents IL_o_ partitioning coefficient for one GPMV. Dotted line represents switch between IL_D_ and IL_O_ phase preference. N = ≥ 8 experiments. **(D)** Summary table for phase preference in the different cell lines. **(E)** GM1 C18:0 (green) and GM1 C18:1^Δ9^ (red) were concomitantly incorporated into the plasma membrane of Hela cells. Scale bar = 1 μm. **(F)** Quantification of GM1 presence in Tfn positive recycling endosomal tubules of Hela cells as for Fig. 1E. Mean ± SEM. **** p = < 0.0001 by One-way ANOVA and Tukey’s multiple comparison. **(G)** Phase preference of different GM1 species, directly fluorescently labeled or clustered by fluorescently labeled CTxB. Dot represents IL_o_ partitioning coefficient of 1 GPMV, n = ≥ 4 experiments. **** p = < 0.0001 by One-way ANOVA and Tukey’s multiple comparison. **(H)** Endosomal sorting of GM1 C16:1^Δ9^ or GM1 C24:1^Δ15^ (green) was monitored in A431 cells in the presence or absence of 100 mM lactose, and transferrin (magenta) and dextran (blue) as in Fig.1C. Scale bars = 5 μm and 0.5 μm respectively. **(I)** FACS sorting of A431 cells incubated with 5 nM fluorescently labeled CTxB in the presence of indicated amount of sucrose or lactose. Cells were then FACS sorted to measure CTxB binding to the cell surface. Lines represent fluorescence of 20,000 cells sorted.

**Suppl. Fig. 2.**
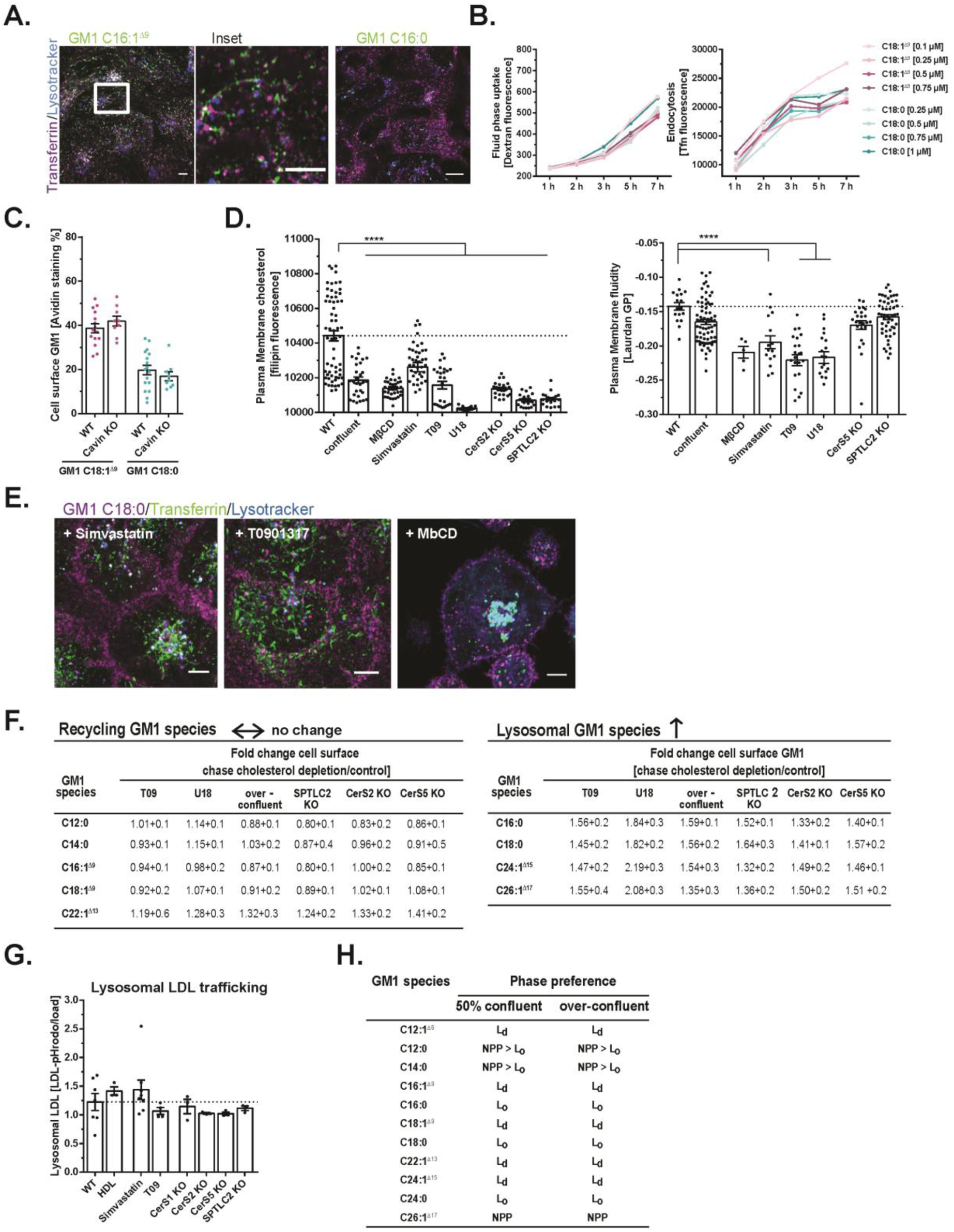
Cholesterol dependent sorting into lysosomal pathway. **(A)** Flu GM1 C16:1^Δ9^ (green) and C16:0 (green) were loaded into A431 WT cells and trafficked with transferrin (magenta) and lysotracker (blue) for 6 h. Recycling endosomal tubules were visualized (Inset). Scale bar = 5 and 1 μm respectively. **(B)** Different concentrations of GM1 C18:1^Δ9^ and GM1 C18:0 were incorporated into A431 cells. GM1 species with C14* motif are displayed in turquoise and GM1 species without C14* motif in magenta. Transferrin and dextran were added during the ‘chase’ period to monitor fluid phase uptake by FACS. Dot represents mean of 2 biological experiments measuring 20,000 cells each. **(C)** GM1 C18:1^Δ9^ and GM1 C18:0 recycling was measured in A431 WT cells or CRISPR-Cas9 KO cells for CAVIN-1 as in (Fig. 1A). Dot represents mean of 2 biological experiments measuring 20,000 cells each. Unpaired t-test was performed. **(D)** A431 WT control cells and cells treated to mildy lower cholesterol were fixed and stained for plasma membrane cholesterol quantification using filipin III (left panel). Cell outlines were detected and mean grey values plotted. Right panel: Membrane fluidity was measured in GPMVs obtained from control and treated cells using laurdan. Each dot represents the general polarization (GP) value for one GPMV. Mean and SEM. **** p = < 0.0001 by One-way ANOVA and Tukey’s multiple comparison. **(E)** GM1 C18:0 (magenta) was incorporated into A431 cells treated as indicated. Tfn (green) and lysotracker™ (blue) were added to demark integrity of intracellular organelles. **(F)** Tabularized summary of plasma membrane recycling for different GM1 species under respective cholesterol lowering conditions as in Fig. 2D. Fold change of GM1 amount at the plasma membrane in treated over control conditions are displayed. ** p = < 0.01 or n.s. by One-way ANOVA and Tukey’s multiple comparison. **(G)** Lysosomal trafficking of pHrodo™-labeled LDL by FACS for A431 cells untreated or treated as indicated (as in D and F). Dot represents mean of 2 biological experiments with 20,000 cells each. Mean ± SEM. One-way ANOVA and Tukey’s multiple comparison. **(H)** Phase preference was quantified different GM1 species in A431 cell GPMVs at 50% confluency or grown to over-confluency for 48 h.

**Suppl. Fig. 3.**
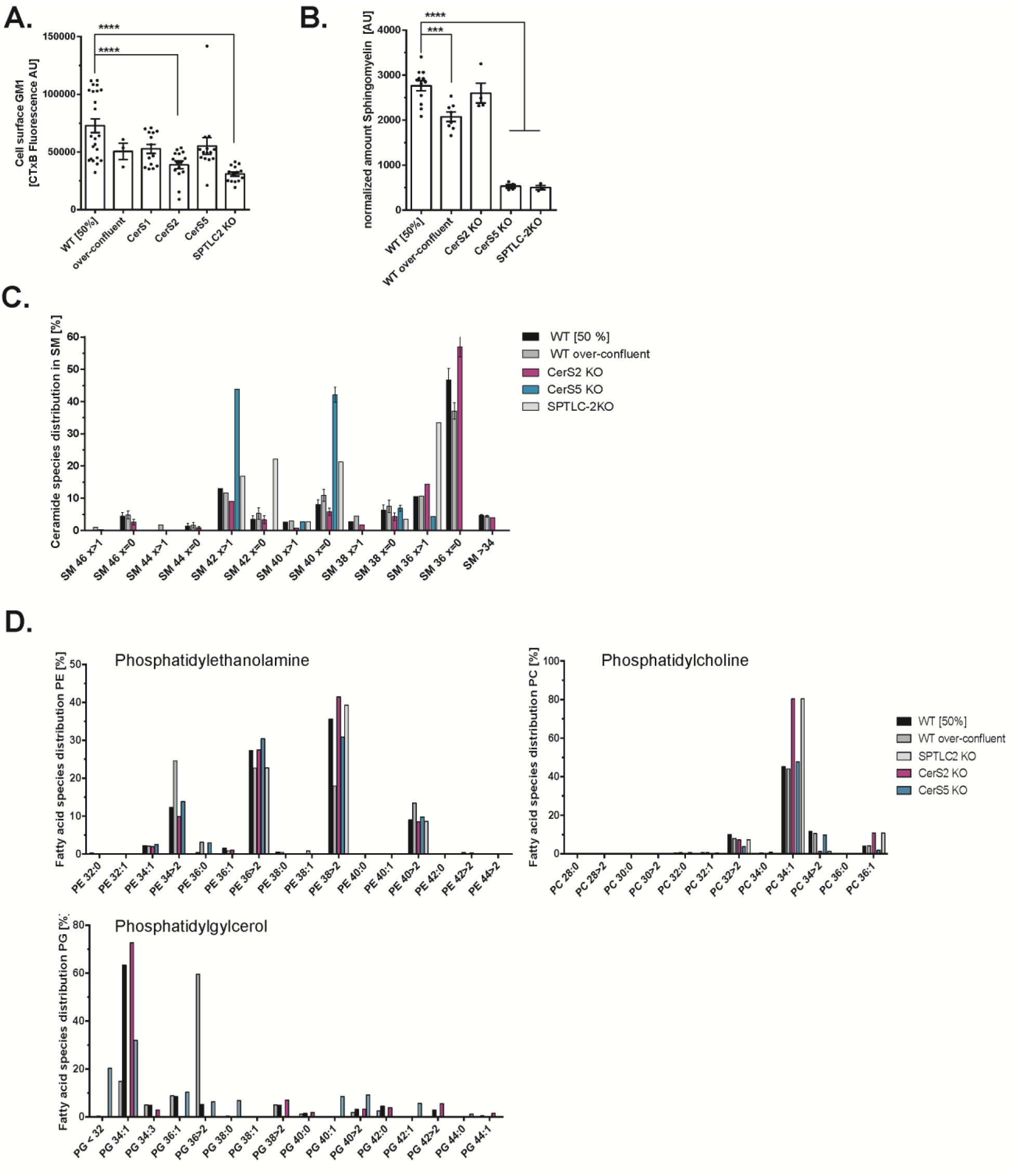
Lipidomic profiling of A431 WT and ceramide synthase KO cell lines. **(A)** CTxB surface staining for GM1 amount in A431 WT or CRISPR-Cas9 KO as indicated. Mean ± SEM. Dot represents mean of 2 biological experiments measuring 20,000 cells each. **** p = < 0.0001 by One-way ANOVA and Tukey’s multiple comparison. **(B)** Sphingomyelin amount of lipidomics profiling for A431 WT, over-confluent and CRISPR-Cas9 KO as indicated. Mean ± SEM. **** p = < 0.0001 or *** p = < 0.001 by One-way ANOVA and Tukey’s multiple comparison. **(C)** Ceramide profile of sphingomyelin species in % for A431 WT, over-confluent and CRISPR-Cas9 KO as indicated. N = > 4 biological replicates. (D) Phosphatidylcholine (PC), - glycerol (PG) and – ethanolamine (PE) fatty acid profiles in % for A431 WT, over-confluent and CRISPR-Cas9 KO as indicated.

**Suppl. Fig. 4.**
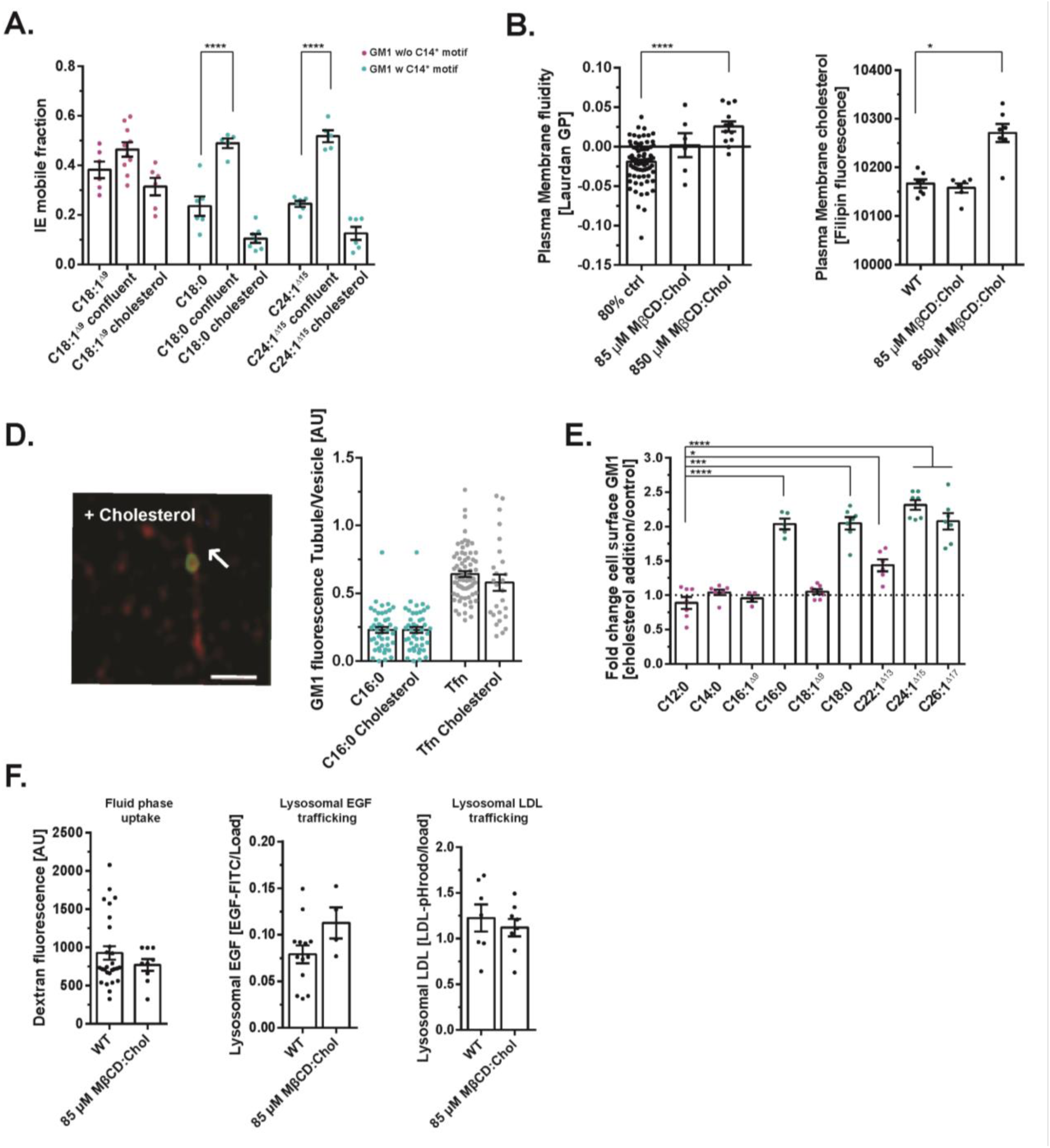
Plasma membrane nanodomain behavior of different GM1 species. **(A)** Quantification of IE mobile fraction for FRAP experiments in Fig. 4C, F and I. Dot represents mean grey fluorescence for one FRAP experiment. Mean ± SEM. **** p = < 0.0001 by One-way ANOVA and Tukey’s multiple comparison. **(B)** Left panel: Filipin staining of plasma membranes of A431 WT cell treated with exogenous cholesterol or untreated. Right panel: Laurdan staining of GPMVs produced from A431 control or cholesterol treated A431 cells Dot represents mean of 2 biological replicates or one GPMV respectively. Mean ± SEM. **** p = < 0.0001 or * p = < 0.05 by One-way ANOVA and Tukey’s multiple comparison. **(D**) Left panel: GM1 C16:0 (green) and transferrin (red) were added to A431 cells treated with exogenous cholesterol. Right panel: Ratio of GM1 C16:0 and Tfn in recycling endosomal tubules over its respective vesicle. Dots represent 1 tubule to vesicle ratio. Mean ± SEM. One-way ANOVA and Tukey’s multiple comparisons were performed. **(E)** FACS based recycling assay for linker-peptide labeled GM1 species in cholesterol treated A431 cells. Fold change between treatment and control is plotted. Dot represents mean of two biological replicates measuring 20,000 cells. Mean ± SEM. **** p = < 0.0001, *** p = < 0.001, * p = < 0.05 by One-way ANOVA and Tukey’s multiple comparison. **(F)** Left panel: dextran fluid phase uptake by FACS was measured in A431 WT cells treated with exogenous cholesterol or untreated. Center and right panel: Lysosomal EGF and LDL trafficking in A431 WT cells treated with exogenous cholesterol or untreated using FITC-EGF and pHrodo™-LDL. Dot represents mean of two biological replicates measuring 20,000 cells. Mean ± SEM. One-way ANOVA and Tukey’s multiple comparisons were performed to test statistical difference. **** p = < 0.0001, *** p = < 0.001, * p = < 0.05 by One-way ANOVA and Tukey’s multiple comparison.

## Supplementary Video Files

**Suppl. Video1.** Lattice lightsheet microscopy of sorting endosome containing GM1 C16:1^Δ9^ (green), transferrin (magenta) and dextran (turquoise) with sorting tubules emerging.

**Suppl. Video1.** Lattice lightsheet microscopy of sorting endosome containing GM1 C18:1^Δ9^ (green), and GM1 C18:0 (magenta) with sorting tubules emerging.

## References

1. M. Koval, R. E. Pagano, Intracellular transport and metabolism of sphingomyelin. Biochimica et biophysica acta 1082, 113 (Mar 12, 1991).

2. G. van Meer, K. Simons, Viruses budding from either the apical or the basolateral plasma membrane domain of MDCK cells have unique phospholipid compositions. The EMBO journal 1, 847 (1982/07/01, 1982).

3. B. Ramstedt, J. P. Slotte, Membrane properties of sphingomyelins. FEBS Lett 531, 33 (Oct 30, 2002).

4. S. Chiantia, E. London, in *Sphingolipids: Basic Science and Drug Development*, E. Gulbins, I. Petrache, Eds. (Springer Vienna, Vienna, 2013), pp. 33–55.

5. D. A. Brown, E. London, Structure and origin of ordered lipid domains in biological membranes. J Membr Biol 164, 103 (Jul 15, 1998).

6. E. London, How principles of domain formation in model membranes may explain ambiguities concerning lipid raft formation in cells. Bba-Mol Cell Res 1746, 203 (Dec 30, 2005).

7. D. Goswami et al., Nanoclusters of GPI-anchored proteins are formed by cortical actin-driven activity. Cell 135, 1085 (Dec 12, 2008).

8. K. Gowrishankar et al., Active remodeling of cortical actin regulates spatiotemporal organization of cell surface molecules. Cell 149, 1353 (Jun 8, 2012).

9. A. Kusumi et al., Membrane mechanisms for signal transduction: the coupling of the meso-scale raft domains to membrane-skeleton-induced compartments and dynamic protein complexes. Seminars in cell & developmental biology 23, 126 (Apr, 2012).

10. C. A. Day, A. K. Kenworthy, Mechanisms underlying the confined diffusion of cholera toxin B-subunit in intact cell membranes. PLoS One 7, e34923 (2012).

11. A. Fujita, J. Cheng, T. Fujimoto, Segregation of GM1 and GM3 clusters in the cell membrane depends on the intact actin cytoskeleton. Biochimica et biophysica acta 1791, 388 (May, 2009).

12. L. Johannes, R. G. Parton, P. Bassereau, S. Mayor, Building endocytic pits without clathrin.

13. L. J. Pike, Lipid rafts: heterogeneity on the high seas. The Biochemical journal 378, 281 (Mar 1, 2004).

14. L. Johannes, R. G. Parton, P. Bassereau, S. Mayor, Building endocytic pits without clathrin. Nat Rev Mol Cell Biol 16, 311 (May, 2015).

15. D. Lingwood, K. Simons, Lipid rafts as a membrane-organizing principle. Science 327, 46 (Jan 1, 2010).

16. K. Jacobson, P. Liu, B. C. Lagerholm, The Lateral Organization and Mobility of Plasma Membrane Components. Cell 177, 806 (May 2, 2019).

17. K. Raghunathan, A. K. Kenworthy, Dynamic pattern generation in cell membranes: Current insights into membrane organization. Biochim Biophys Acta Biomembr 1860, 2018 (Oct, 2018).

18. E. Sezgin, I. Levental, S. Mayor, C. Eggeling, The mystery of membrane organization: composition, regulation and roles of lipid rafts. Nat Rev Mol Cell Biol 18, 361 (Jun, 2017).

19. K. Simons, J. L. Sampaio, Membrane organization and lipid rafts. Cold Spring Harb Perspect Biol 3, a004697 (Oct 1, 2011).

20. K. Simons, G. van Meer, Lipid sorting in epithelial cells. Biochemistry 27, 6197 (1988).

21. P. Varshney, V. Yadav, N. Saini, Lipid rafts in immune signalling: current progress and future perspective. Immunology 149, 13 (Sep, 2016).

22. Y. Zhou, J. F. Hancock, Ras nanoclusters: Versatile lipid-based signaling platforms. Biochimica et biophysica acta 1853, 841 (Apr, 2015).

23. M. F. Garcia-Parajo, A. Cambi, J. A. Torreno-Pina, N. Thompson, K. Jacobson, Nanoclustering as a dominant feature of plasma membrane organization. Journal of cell science 127, 4995 (2014).

24. J. Lippincott-Schwartz, R. D. Phair, Lipids and cholesterol as regulators of traffic in the endomembrane system. Annu Rev Biophys 39, 559 (2010).

25. G. H. Patterson et al., Transport through the Golgi apparatus by rapid partitioning within a two-phase membrane system. Cell 133, 1055 (Jun 13, 2008).

26. Y. Deng, F. E. Rivera-Molina, D. K. Toomre, C. G. Burd, Sphingomyelin is sorted at the trans Golgi network into a distinct class of secretory vesicle. Proc Natl Acad Sci U S A 113, 6677 (Jun 14, 2016).

27. M. B. Stone, S. A. Shelby, M. F. Nunez, K. Wisser, S. L. Veatch, Protein sorting by lipid phase-like domains supports emergent signaling function in B lymphocyte plasma membranes. Elife 6, e19891 (Feb 1, 2017).

28. W. I. Lencer, B. Tsai, The intracellular voyage of cholera toxin: going retro. Trends in biochemical sciences 28, 639 (Dec, 2003).

29. P. Cuatrecasas, Gangliosides and membrane receptors for cholera toxin. Biochemistry 12, 3558 (Aug 28, 1973).

30. M. A. Campanero-Rhodes et al., N-glycolyl GM1 ganglioside as a receptor for simian virus 40. J Virol 81, 12846 (Dec, 2007).

31. K. Sandvig, Shiga toxins. Toxicon 39, 1629 (Nov, 2001).

32. M. Lorizate et al., Comparative lipidomics analysis of HIV-1 particles and their producer cell membrane in different cell lines. Cell Microbiol 15, 292 (Feb, 2013).

33. W. Romer et al., Shiga toxin induces tubular membrane invaginations for its uptake into cells. Nature 450, 670 (Nov 29, 2007).

34. R. Lakshminarayan et al., Galectin-3 drives glycosphingolipid-dependent biogenesis of clathrin-independent carriers. Nat Cell Biol 16, 595 (Jun, 2014).

35. Y. A. Hannun, L. M. Obeid, Sphingolipids and their metabolism in physiology and disease. Nat Rev Mol Cell Biol 19, 175 (Mar, 2018).

36. A. H. Merrill, Jr., Sphingolipid and glycosphingolipid metabolic pathways in the era of sphingolipidomics. Chem Rev 111, 6387 (Oct 12, 2011).

37. W. W. Young, M. S. Lutz, W. A. Blackburn, Endogenous glycosphingolipids move to the cell surface at a rate consistent with bulk flow estimates. Journal of Biological Chemistry 267, 12011 (1992/06/15/, 1992).

38. G. van Meer, D. R. Voelker, G. W. Feigenson, Membrane lipids: where they are and how they behave. Nat Rev Mol Cell Biol 9, 112 (Feb, 2008).

39. D. E. Saslowsky et al., Ganglioside GM1-mediated transcytosis of cholera toxin bypasses the retrograde pathway and depends on the structure of the ceramide domain. J Biol Chem 288, 25804 (Sep 6, 2013).

40. D. J. Chinnapen et al., Lipid sorting by ceramide structure from plasma membrane to ER for the cholera toxin receptor ganglioside GM1. Developmental cell 23, 573 (Sep 11, 2012).

41. S. Rodriguez-Gallardo et al., Ceramide chain length-dependent protein sorting into selective endoplasmic reticulum exit sites. Sci Adv 6, (Dec, 2020).

42. C. M. Paton, J. M. Ntambi, Biochemical and physiological function of stearoyl-CoA desaturase. American journal of physiology. Endocrinology and metabolism 297, E28 (Jul, 2009).

43. B. Ramstedt, J. P. Slotte, Interaction of cholesterol with sphingomyelins and acyl-chain-matched phosphatidylcholines: a comparative study of the effect of the chain length. Biophys J 76, 908 (Feb, 1999).

44. A. Kihara, Very long-chain fatty acids: elongation, physiology and related disorders. Journal of biochemistry 152, 387 (Nov, 2012).

45. S. Kavaliauskiene, et al., Cell density-induced changes in lipid composition and intracellular trafficking. Cellular and molecular life sciences : CMLS 71, 1097 (Mar, 2014).

46. R. L. Shaner et al., Quantitative analysis of sphingolipids for lipidomics using triple quadrupole and quadrupole linear ion trap mass spectrometers. J Lipid Res 50, 1692 (Aug, 2009).

47. Y. M. te Welscher, D. J. Chinnapen, L. Kaoutzani, R. J. Mrsny, W. I. Lencer, Unsaturated glycoceramides as molecular carriers for mucosal drug delivery of GLP-1. J Control Release 175, 72 (Feb 10, 2014).

48. M. D. Garcia-Castillo et al., Mucosal absorption of therapeutic peptides by harnessing the endogenous sorting of glycosphingolipids. Elife 7, e34469 (May 31, 2018).

49. S. L. Veatch, S. L. Keller, Separation of liquid phases in giant vesicles of ternary mixtures of phospholipids and cholesterol. Biophys J 85, 3074 (Nov, 2003).

50. E. Sezgin et al., Elucidating membrane structure and protein behavior using giant plasma membrane vesicles. Nat Protoc 7, 1042 (May 3, 2012).

51. F. R. Maxfield, T. E. McGraw, Endocytic recycling. Nat Rev Mol Cell Biol 5, 121 (Feb, 2004).

52. S. Mukherjee, R. N. Ghosh, F. R. Maxfield, Endocytosis. Physiol Rev 77, 759 (Jul, 1997).

53. S. B. Kellie, B. Patel, J. Pierce, D. Critchley, Capping of cholera toxin - Ganglioside GM1 complexes on mouse lymphocytes is accompanied by co-capping of alpha-actinin. J. Cell Biol. 97, 447 (1983).

54. A. M. Kabbani, K. Raghunathan, W. I. Lencer, A. K. Kenworthy, C. V. Kelly, Structured clustering of the glycosphingolipid GM1 is required for membrane curvature induced by cholera toxin. Proceedings Of The National Academy Of Sciences Of The United States Of America. 117, 14978 (Jun 30, 2020).

55. B. Sorre et al., Curvature-driven lipid sorting needs proximity to a demixing point and is aided by proteins. Proc Natl Acad Sci U S A 106, 5622 (Apr 7, 2009).

56. B. N. Stillman et al., Galectin-3 and galectin-1 bind distinct cell surface glycoprotein receptors to induce T cell death. Journal of immunology 176, 778 (Jan 15, 2006).

57. S. Mayor, J. F. Presley, F. R. Maxfield, Sorting of membrane components from endosomes and subsequent recycling to the cell surface occurs by a bulk flow process. J Cell Biol 121, 1257 (Jun, 1993).

58. A. A. Wolf et al., Attenuated Endocytosis and Toxicity of a Mutant Cholera Toxin with Decreased Ability to Cluster Gm1. Infect Immun 76, 1476 (Jan 22, 2008).

59. K. Raghunathan et al., Glycolipid Crosslinking Is Required for Cholera Toxin to Partition Into and Stabilize Ordered Domains. Biophys J, (Nov 30, 2016).

60. P. Luong et al., A quantitative single-cell assay for retrograde membrane traffic enables rapid detection of defects in cellular organization. Molecular biology of the cell 31, 511 (2020).

61. J. W. Park, W. J. Park, A. H. Futerman, Ceramide synthases as potential targets for therapeutic intervention in human diseases. Biochimica et biophysica acta 1841, 671 (May, 2014).

62. K. M. Spillane et al., High-speed single-particle tracking of GM1 in model membranes reveals anomalous diffusion due to interleaflet coupling and molecular pinning. Nano Lett 14, 5390 (Sep 10, 2014).

63. J. S. Goodwin, K. R. Drake, C. L. Remmert, A. K. Kenworthy, Ras diffusion is sensitive to plasma membrane viscosity. Biophys J 89, 1398 (Aug, 2005).

64. A. Roux et al., Role of curvature and phase transition in lipid sorting and fission of membrane tubules. The EMBO journal 24, 1537 (Apr 20, 2005).

65. T. Baumgart, S. T. Hess, W. W. Webb, Imaging coexisting fluid domains in biomembrane models coupling curvature and line tension. Nature 425, 821 (Oct 23, 2003).

66. S. Mukherjee, T. T. Soe, F. R. Maxfield, Endocytic sorting of lipid analogues differing solely in the chemistry of their hydrophobic tails. J Cell Biol 144, 1271 (Mar 22, 1999).

67. P. Noutsi, E. Gratton, S. Chaieb, Assessment of Membrane Fluidity Fluctuations during Cellular Development Reveals Time and Cell Type Specificity. PLoS One 11, e0158313 (2016).

68. J. Steinkuhler, E. Sezgin, I. Urbancic, C. Eggeling, R. Dimova, Mechanical properties of plasma membrane vesicles correlate with lipid order, viscosity and cell density. Commun Biol 2, 337 (2019/09/13, 2019).

69. M. Hao, S. Mukherjee, Y. Sun, F. R. Maxfield, Effects of cholesterol depletion and increased lipid unsaturation on the properties of endocytic membranes. The Journal of biological chemistry 279, 14171 (Apr 2, 2004).

70. A. Kihara, Very long-chain fatty acids: elongation, physiology and related disorders.

71. M. L. Fanani, B. Maggio, The many faces (and phases) of ceramide and sphingomyelin I - single lipids. Biophys Rev 9, 589 (Oct, 2017).

72. M. L. Fanani, B. Maggio, The many faces (and phases) of ceramide and sphingomyelin II - binary mixtures. Biophys Rev 9, 601 (Oct, 2017).

73. G. W. Stockton, I. C. Smith, A deuterium nuclear magnetic resonance study of the condensing effect of cholesterol on egg phosphatidylcholine bilayer membranes. I. Perdeuterated fatty acid probes. Chem Phys Lipids 17, 251 (Oct, 1976).

74. S. Jaikishan, A. Bjorkbom, J. P. Slotte, Sphingomyelin analogs with branched N-acyl chains: the position of branching dramatically affects acyl chain order and sterol interactions in bilayer membranes. Biochimica et biophysica acta 1798, 1987 (Oct, 2010).

75. K. Simons, W. L. Vaz, Model systems, lipid rafts, and cell membranes. Annu Rev Biophys Biomol Struct 33, 269 (2004/06/09, 2004).

76. R. Raghupathy et al., Transbilayer lipid interactions mediate nanoclustering of lipid-anchored proteins. Cell 161, 581 (Apr 23, 2015).

77. T. Fujimoto, I. Parmryd, Interleaflet Coupling, Pinning, and Leaflet Asymmetry—Major Players in Plasma Membrane Nanodomain Formation. Frontiers in Cell and Developmental Biology 4, 155 (2017).

78. T. Rog et al., Interdigitation of long-chain sphingomyelin induces coupling of membrane leaflets in a cholesterol dependent manner. Biochimica et biophysica acta 1858, 281 (Feb, 2016).

79. T. Skotland, K. Sandvig, The role of PS 18:0/18:1 in membrane function. Nat Commun 10, 2752 (Jun 21, 2019).

80. T. K. M. Nyholm et al., Impact of Acyl Chain Mismatch on the Formation and Properties of Sphingomyelin-Cholesterol Domains. Biophys J 117, 1577 (Nov 5, 2019).

81. Y. Y. Chou et al., Identification and Characterization of a Novel Broad-Spectrum Virus Entry Inhibitor. J Virol 90, 4494 (May, 2016).

82. M. Nissim-Rafinia, E. Meshorer, Photobleaching assays (FRAP & FLIP) to measure chromatin protein dynamics in living embryonic stem cells. Journal of visualized experiments : JoVE, e2696 (Jun 29, 2011).

83. L. Liaw et al., Lipid Profiling of In Vitro Cell Models of Adipogenic Differentiation: Relationships With Mouse Adipose Tissues. J Cell Biochem 117, 2182 (Sep, 2016).

84. N. Gajenthra Kumar et al., Untargeted lipidomic analysis to broadly characterize the effects of pathogenic and non-pathogenic staphylococci on mammalian lipids. PLoS One 13, e0206606 (2018).

85. J. G. Doench et al., Optimized sgRNA design to maximize activity and minimize off-target effects of CRISPR-Cas9. Nat Biotechnol 34, 184 (Feb, 2016).

